# Nuclei sense complex tissue shape and direct intestinal stem cell fate

**DOI:** 10.1101/2025.10.23.684219

**Authors:** Kaustav Bera, Delaney L. McNally, Bruce E. Kirkpatrick, Nolan R. Petrich, F. Max Yavitt, Marilyne Coulombe, Michaela Quintero, Nathaniel P. Skillin, Alex Khang, Patrick S. McGrath, Linda C. Samuelson, Tanmay P. Lele, Peter J. Dempsey, Kristi S. Anseth

## Abstract

Tissue architecture and function are influenced by mechanical cues. Yet, how cell nuclei sense forces within 3D tissues and dictate differentiation remains unknown as prior studies focused on isolated mesenchymal cells, which fail to fully predict tissue-level mechanical properties. We fill this knowledge gap utilizing live reporters and material-based organoid models. We posit the nucleus as an active mechanosensor of tissue shape, with levels of the nuclear scaffolding protein lamin-A varying across intestinal stem cell differentiation trajectory. Elevated forces on differentiated Paneth cell nuclei, in both organoids and tissue explants, increase lamin-A and nuclear wrinkling. Enhancing nuclear mechanotransduction primes cell differentiation, in otherwise stem promoting conditions, revealing that nuclear mechanics can direct stem cell fate. By engineering spatiotemporally controlled *de novo* tissue curvature with photo-degradable hydrogels, we direct spatially patterned lamin-A levels across mouse and human organoids of healthy and diseased origin, uncovering conserved nuclear mechanosensing pathway in epithelial tissues.

## Introduction

Epithelial tissues are thin layers of tightly packed cells that actively respond to mechanical forces while carrying out a variety of functions like maintaining organ shape and providing chemical and mechanical barriers^1^. During development, forces on the epithelia from the surrounding microenvironment sculpt tissue architecture - driving processes such as villus morphogenesis in the intestine and airway branching in the lung^2^. In addition to terminally differentiated cells, a majority of adult epithelial tissues maintain spatially defined stem cell populations to replenish cells lost during healthy organ functioning or injury^3^. Due to their mixed composition and continuous cell fate transitions, epithelial tissues behave as complex living materials with spatiotemporally variant mechanical properties^4,5^.

Understanding how physical forces and niche mechanics regulate epithelial cell behavior is essential for predicting tissue homeostasis and intervening in diseases characterized by epithelial dysregulation. While it is known that the nucleus in a variety of cells confers cellular mechanosensitivity by integrating mechanical signals, these insights are largely based on studies of single mesenchymal cells interacting with a matrix^6,7^. In contrast cells in epithelial tissues are densely packed with many cell-cell contacts, and how cells in these three-dimensional (3D) microenvironment sense and respond to mechanical cues remains unknown.

The nucleus is the stiffest organelle in the cell and harbors its genetic material surrounded by a double membrane nuclear envelope (NE). The NE is supported underneath by a mesh-like nuclear lamina composed of intermediate filaments of A or C and B type lamins^7^. Application of mechanical forces on the nuclei, for example through cellular exposure to stiff microenvironment or compression, can restructure the NE by elevating lamin A levels^8^ and opening nuclear pore complexes^9^ that facilitate nuclear entry of mechanosensitive transcriptional regulators such as Yes-associated protein (YAP). Given the paucity of information, we sought to elucidate how cell nuclei embedded in a 3D epithelium would respond to tissue shape changes and what role these changes would play in regulating cell fate during stem differentiation.

With this in mind, the inner lining of the mammalian intestine provides a robust system to study the role of nuclear sensing in 3D tissues, as the intestinal epithelium has a highly defined yet complex tissue architecture^10^. The intestinal epithelium is arranged in a crypt-villus array which is composed of multiple cell types responsible for the tissue’s specialized functions (**Fig. 1a**). The crypt base contains multipotent Leucine-rich repeat-containing G protein-coupled receptor 5 positive (LGR5+) intestinal stem cells (ISCs) that rapidly divide to replace differentiated epithelial cells, the majority of which are shed off every 5-7 days due to the presence of constant chemical and physical stressors^11,12^. Within the crypt, ISCs co-exist with differentiated Paneth cells that are responsible for secreting lysozyme and maintaining the stem cell niche within the crypt. Localized to the crypt niche are key biochemical factors like WNT, NOTCH and bone morphogenetic protein (BMP) that tightly regulate ISC proliferation, self-renewal and differentiation^13^. In addition to biochemical cues, the basement membrane and its mechanical properties can regulate crypt cell composition through the mechanosensitive PIEZO ion channels^14^. 3D organoids derived from ISCs have been grown *in vitro* and preserve this mechanoresponsiveness, where their differentiation and cellular composition can be controlled by engineered epithelial tissue geometries that alter ion channel activation and YAP signaling^15–18^.

Result herein demonstrate that Paneth cells in mouse intestinal tissues and organoids have different nuclear shape and organization compared to non-Paneth cells. Notably, the Paneth cell NE displays elevated deformation and lamin A, indicative of higher forces that were further verified by laser ablation studies. Encapsulating spherical organoids in photo-degradable materials allowed control over symmetry breaking and *de novo* bud formation, revealing that changes in the NE occur rapidly (≤3 hours) after epithelial deformation. At this short timescale, YAP is not yet upregulated, indicating a direct biophysical response of the nuclei in response to tissue buckling. Increasing mechanosensitivity of the cell nuclei by enriching lamin A at the NE further promotes ISC differentiation independent of biochemical cues presented in the medium. Taken together, our findings reveal that cell nuclei in epithelial tissues function as active mechanosensors, translating physical cues into cell fate decisions. Then, by photo-patterning human derived organoids, we confirm that this nuclear mechanosensing is preserved across a variety of epithelial tissues.

## Results

### Differentiated Paneth cells display distinct nuclear organization, shape, and composition both *in vivo* and *in vitro*

To understand the mechanical state of cell nuclei during epithelial homeostasis in intestinal crypts, we analyzed the shape, location, and orientation of nuclei in crypts within the mouse small intestine (**Fig. 1b**). We observed that the nuclei of lysozyme-positive Paneth cells were more circular compared to non-Paneth cell nuclei which were more elongated. The nuclear shapes were quantified by measuring the aspect ratio, which was lower in Paneth compared to non-Paneth cells (**Fig. 1c**). Beyond this difference, Paneth cell nuclei were also located closer and aligned parallel to the basal surface of the epithelia, as quantified by the shortest distance of the nuclear centroid and orientation of the nuclear major axis, respectively (**Fig. 1b, d-f**). These *in vivo* crypt observations were complemented with *in vitro* experiments using murine jejunal ISC-derived organoid cultures in Matrigel (**Extended Fig. 1a, Fig. 1g**), which grow crypt-like buds under differentiation conditions (ENR medium). Analysis of organoid crypt buds also revealed a smaller aspect ratio, shorter distance, and closer to parallel alignment of Paneth cell nuclei to the epithelial base (**Fig. 1c-f**). These findings suggested a conserved nuclear morphology characteristic of differentiated Paneth cells as compared to non-differentiated crypt stem and progenitor cells.

Differences in the arrangement and morphology of Paneth cell nuclei suggested that they might be subject to elevated mechanical compression from the cytoplasmic lysozyme granules, prompting us to examine the forces experienced by nuclei within the epithelium. To investigate this, we first looked at the levels of lamin A in the nuclei of organoid crypts using immunofluorescence (**Fig. 1h**). Lamin A levels have been shown to scale directly with the amount of force a cell nucleus experiences^8,19^. In line with our earlier observations, Paneth cell nuclei expressed high levels of lamin A, suggesting higher force upon the nuclear envelope (**Fig. 1i**). Conversely, if organoids were kept in stem promoting conditions (ENRCV growth medium between days zero to three in the timeline from **Extended Fig. 1a)**, the growing epithelial enteroids did not produce lysozyme-positive Paneth cells (or a scarce few), and the nuclei had a nearly homogeneous distribution of lamin A levels throughout the organoid (**Extended Fig. 1b**). To obtain an unbiased measure of the nuclear mechanical state, we co-stained organoids derived from Paneth cell reporter mice (*Defa4^Cre^/Rosa26^tdTomato^*)^15,20^ with antibodies against lamin A and lamin B (**Fig. 1j**). *Defa4^Cre^*-expressing Paneth cells could be identified from tdTomato expression and their nuclei showed slightly lower levels of lamin B (**Fig. 1j, k**) while their lamin A levels were elevated, supporting prior observations (**Fig. 1j, Extended Fig. 1c**). Interestingly, a pixel-based ratio of lamin A over lamin B levels distinctly demarcated the Paneth cell nuclei (**Fig. 1j**, **l**).

As an orthogonal approach to identify the forces upon the nuclei, we examined wrinkling^6,21,22^ within the nuclei in organoid crypts. Staining of both DNA and the nuclear lamina (lamin A) identified high degrees of nuclei deformation, but only in the Paneth cells (**Fig. 1m**). Near super resolution imaging of cell nuclei in the 3D organoids was achieved using photo-expansion microscopy (PhotoExM)^23,24^, which further revealed furrows where the NE protruded into the DAPI stained DNA during wrinkling (**Extended Fig. 1d, e**). Cumulatively, this data identifies key differences in the nuclear shape and lamina composition in intestinal crypt cells (i.e., the stem cell niche), hinting towards unequal forces acting upon differentiated Paneth cells as compared to ISCs and other non-Paneth cells that reside in the crypt.

**Figure 1:**
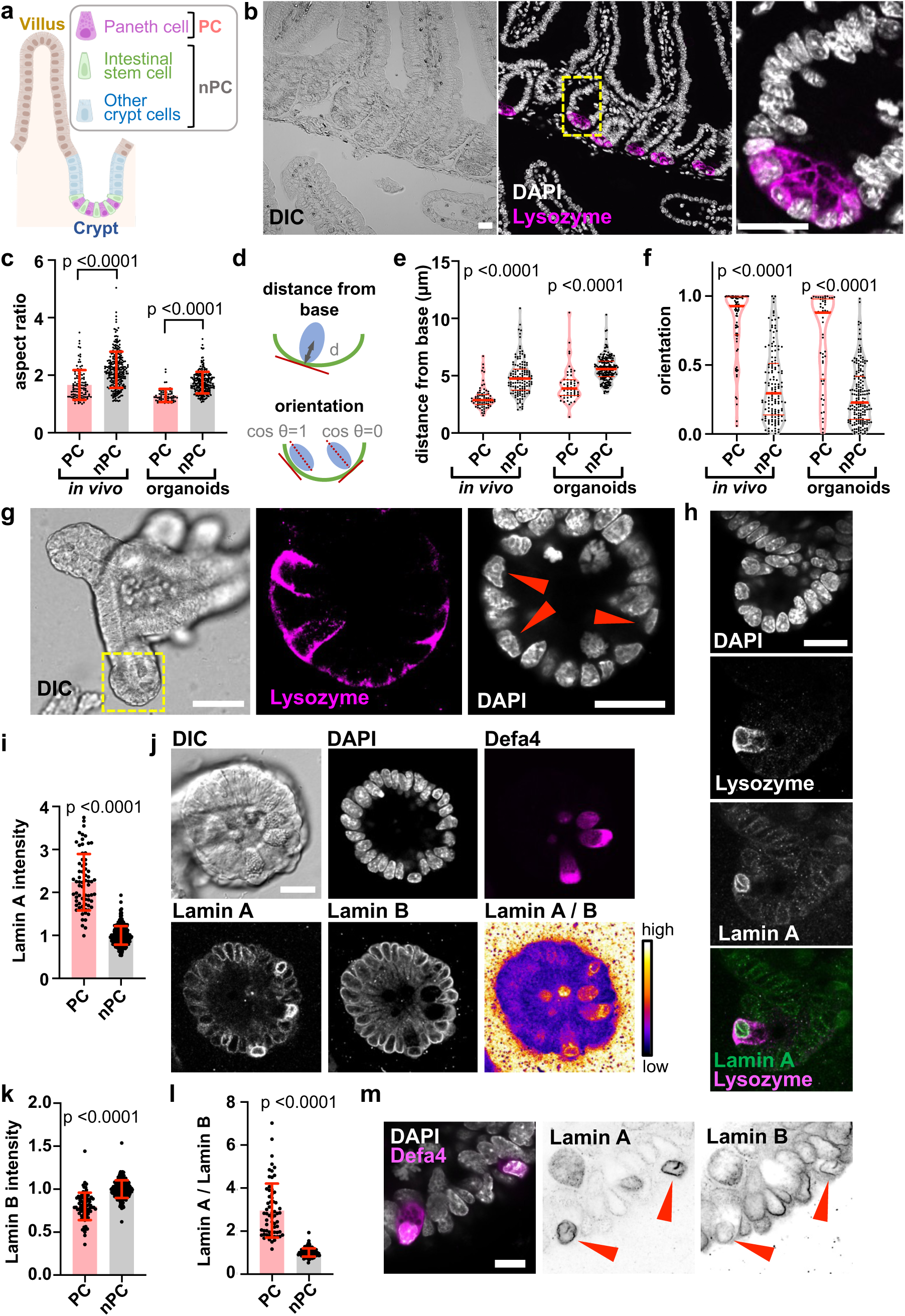
Differentiated Paneth cells in the mouse intestine and *in vitro* organoids have distinct nuclear arrangement, shape, and composition. **a**, A schematic crypt-villus unit of the intestinal epithelium with intestinal stem cells (ISCs) and lysozyme+ Paneth cells. Cells in brown and blue represent other cells from the villus and crypt, respectively. For subsequent studies, crypt cells have been categorized into Paneth cells (PC) or non-Paneth cells (nPC). **b,** Confocal images of lysozyme and DAPI stained mouse intestinal sections. Image at the right shows magnified region indicated with the yellow rectangle in the image at the center. Images are representative of 3 mice. Scale bar = 20 µm **c**, Aspect ratio of nuclei in Paneth cells and non-Paneth cells in mouse intestine and murine ISC derived organoids grown and differentiated in Matrigel. Data are mean ± s.d. from 3 mice and 3 *in vitro* experiments. **d**, *Top*, distance from base is calculated by finding the shortest distance between the centroid of the nucleus and the basal surface of the epithelium. *Bottom*, orientation of the nucleus is reported as the cosine of the angle between the major axis of the nuclei and the tangent to the epithelial base at the point closest to the centroid of the nucleus. **e**, **f**, Distance of nuclei from epithelium base (**e**) and orientation of nuclei (**f**) in mouse tissue sections (from 3 mice) and ISC derived organoids from 3 experiments. Red lines indicate the median (thick) and quartiles (thin). **g**, *Left*, confocal DIC image of differentiated organoid (scale bar = 50 µm), *Center* and *Right*, confocal images of lysozyme and DAPI stained organoid bud enclosed by yellow box in DIC image (scale bar = 20 µm). Red arrowheads indicate Paneth cell nuclei. Images are representative of 3 experiments. **h**, Immuno-stained confocal images of organoid buds. Images are representative of 3 experiments. Scale bar = 20 µm. **i**, Lamin A intensities of nuclear periphery in Paneth and non-Paneth cells within organoid buds. Values are normalized to average of non-Paneth cell intensities in each image. Data are mean ± s.d. from 3 experiments. **j**, Immuno-stained confocal images of *Defa4^Cre^*/*Rosa26^tdTomato^*organoid buds. Heatmap show pixelwise ratio of lamin A over lamin B immunofluorescence. Images are representative of 3 experiments. Scale bar = 20 µm. **k**, Lamin B intensities of nuclear periphery in Paneth and non-Paneth cells within organoid buds. Values are normalized to average of nPC intensities in each image. Data are mean ± s.d. from 3 experiments. **l**, Ratio of lamin A over lamin B immunofluorescence intensities in nuclear lamina of Paneth and non-Paneth cells. Data are mean ± s.d. from 3 experiments. **m**, Confocal image of DAPI, lamin A, and lamin B-stained *Defa4^Cre^*/Rosa26*^tdTomato^* organoid bud showing invagination of nuclear lamina during wrinkling in Paneth cells (red arrowheads). Images are representative of 3 experiments. Scale bar = 10 µm. Statistical analysis was performed using Mann-Whitney U-test (for comparing organoids in **c** and **k**, *in vivo* crypts in **c**, **e** and both in **f**) and unpaired t-tests after log-transformation (for comparing organoids in **e**, **i**, and **l**).

### Nuclear deformation in Paneth cells is caused by cytoplasmic forces

Cell nuclei are mechanically linked to the extracellular environment via a cytoskeleton network composed of actin microfilaments. To probe if a reduction in cytoskeletal tension could relieve the force upon the nuclei, we treated *Defa4^Cre^*/Rosa26*^tdTomato^* organoids containing Paneth cells with the F-actin inhibitor Cytochalasin D (**Fig. 2a**). Upon disruption of the actin network, we observed larger changes in the solidity of the Paneth cell nuclei (three times compared to non-Paneth cells), indicative of unfolding of the nuclear wrinkles (**Fig. 2b**). Using live cell tracking, we quantified changes in the cross-sectional area of each nucleus as cytoskeletal forces were released after inhibition of F-actin polymerization (**Extended Fig. 2a**). Results showed an overall increase in the cross-sectional area of the Paneth cell nuclei upon treatment, while the average cross-sectional area of non-Paneth cell nuclei remained unchanged (**Fig. 2c**).

Finally, as a direct demonstration of how the epithelial forces differentially cause a high degree of Paneth cell nuclear deformation, we used laser assisted ablation of cells in a differentiated organoid and tracked ensuing changes in the nucleus. Specifically, *Defa4^Cre^*/Rosa26*^tdTomato^* intestinal organoids with either a live nuclear H2B reporter or Hoechst 33342 staining were differentiated to obtain tdTomato-positive Paneth cells, which were then photo-ablated on the apical cortex of cells using an 800 nm laser to release their cytoplasmic lysozyme granules and thus the force on the nuclei (**Fig. 2d**). Similar to results with bulk inhibition of F-actin (i.e., Cytochalasin D treatment), spatial release of cytoplasmic forces on the nuclei by ablation of the cell cortex increased the nuclear cross-sectional area, specifically in the targeted Paneth cells (**Fig. 2e**, **Extended Fig. 2b**). In contrast, the average cross-sectional area of non-Paneth cell nuclei remained unchanged after apical ablation (**Fig. 2d**, **e**, **Extended Fig. 2b**). These observations cumulatively support the notion that nuclei in Paneth cells experience elevated levels of forces compared to their neighboring non-Paneth cells, altering their nuclear morphology and enriching lamin A in their NE. However, since the Paneth cell cytoplasm is filled with lysozyme granules generating compressive forces on the nucleus, we next wanted to check if elevated lamin A levels arise in nuclei of cells prior to terminal differentiation and lysozyme accumulation.

### Changes in nuclei are manifested in secretory precursor cells

The organoids studied thus far were differentiated for three days (**Extended Fig. 1a**) to obtain terminal Paneth cells within crypts, similar to the *in vivo* intestinal lining. To probe if changes in nuclear lamina primarily exist as a result of lysozyme granule accumulation in the cytoplasm of Paneth cells versus curvature induced forces in the epithelium, we examined lamin A levels in nuclei of secretory precursor cells prior to complete differentiation and lysozyme expression. Delta-like 4 and Delta-like 1 (DLL4 and DLL1) have been used to identify such precursor cells in the intestinal epithelium^18,25^. Thus, using organoids expanded from transgenic *Dll4-mCherry* reporter mice^25^, we verified that the mCherry signal appears within a day after initiating differentiation medium conditions (ENR medium) but prior to detectable lysozyme expression (**Fig. 2f**, **Extended Fig. 2c**). With this reporter organoid system, we observed even at early differentiation timepoints (within a day of ENR medium treatment), mCherry+ precursor secretory cells that showed high levels of lamin A expression, indicating higher nuclear forces (**Fig. 2f, g**). This observation was not limited to the DLL4 reporter organoid line, as we also observed a positive correlation of the lamin A intensity with DLL1 immunostaining in wild-type organoids (**Extended Fig. 2d**). Taken together, these results suggest that production of lysozyme granules in the cytoplasm is not necessary for higher lamin A expression in Paneth cells, and DLL4 or DLL1 positive secretory precursor cells are perhaps primed for differentiation by higher nuclear mechanotransduction. Since the secretory precursor cells showed higher expression of lamin A compared to other cells in the epithelium, we next asked the question when their nuclei begin to sense higher forces during the crypt formation process.

**Figure 2:**
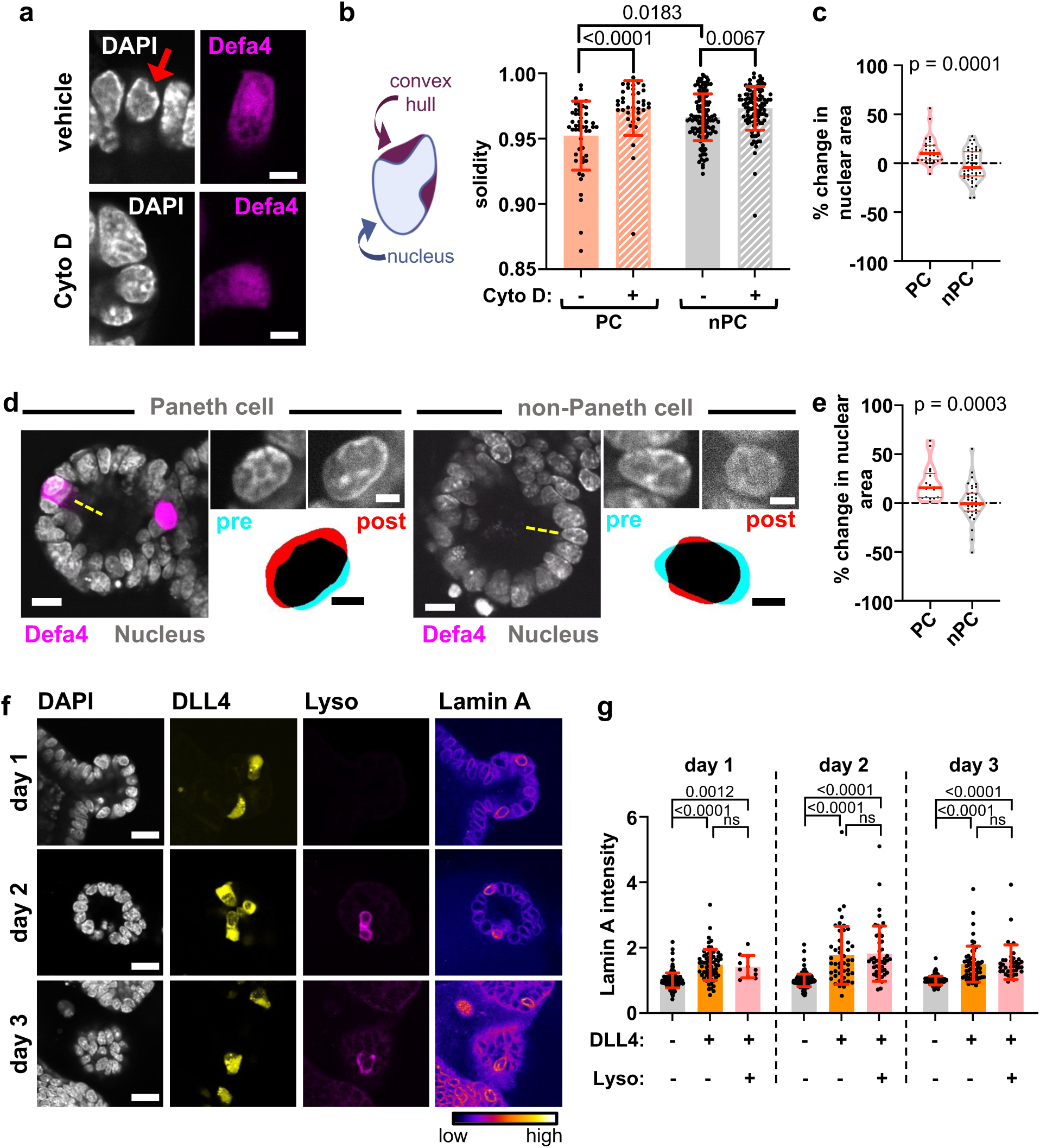
Changes in cell nuclei arise due to spatially variant forces in cells that appear in secretory precursor cells before cytosolic lysozyme accumulation. **a**, Confocal images of DAPI stained *Defa4^Cre^*/*Rosa26^tdTomato^*organoid buds after Cytochalasin D or vehicle control treatment. Red arrow shows nuclear deformation in a *Defa4^Cre^* expressing cell. Images are representative of 3 experiments. Scale bars = 5 µm. b, *Left*, cartoon of the convex hull surrounding a cell nucleus to calculate the solidity (ratio of actual area to that of convex hull). *Right*, Solidity of Paneth (PC) and non-Paneth cells (nPC) nuclei immuno-stained after treatment with Cytochalasin D or vehicle control for 1 hour. Data are mean ± s.d. from 3 experiments. **c**, Percentage change in cross sectional area of Paneth cell and non-Paneth cell nuclei after treatment with Cytochalasin D for 1 hour. The red lines indicate the medians (thick) and quartiles (thin) from 3 experiments. **d**, Representative 2 photon images of *Defa4^Cre^*/*Rosa26^tdTomato^*organoid buds stained for nuclei with Hoechst 33342. Yellow dashed lines represent the regions of laser ablation and scale bars = 10 µm. Insets show nuclei pre- and post-laser ablation of apical cortex of cell (Scale bars = 3 µm). Overlap of cyan (pre) and red (post) nuclei outlines indicate the relative changes in cross sectional area. Images are representative of >3 experiments. **e**, Percentage change in cross sectional area of Paneth cell and non-Paneth cell nuclei after laser ablation from >3 experiments. The red lines indicate the median (thick) and quartiles (thin). **f**, Immuno-stained confocal images of *Dll4-mCherry* expressing organoids stained for lamin A and lysozyme at indicated timepoints after switching medium to differentiation (ENR) conditions. Images are representative of 3 experiments and scale bars = 20 µm. **g**, Lamin A intensities of NE in cell populations segregated based upon their lysozyme and DLL4 expression. Data are mean ± s.d. from 3 experiments. Statistical analysis was performed using Kruskal Wallis test followed by Dunn’s test (in **b** and **g**) and Mann-Whitney U-test (in **c** and **e**).

### Photo-degradable hydrogels allow spatiotemporal control of mechanical force evolution and crypt budding

To elucidate the time resolved dynamics of the nuclear lamina, we exploited a spatiotemporally controllable crypt formation model^15,18^. Poly(ethylene glycol) (PEG) hydrogels were prepared at 2.5 wt% by reacting 8-arm 40 kDa PEG star macromolecules functionalized with nitrobenzyl azide or dibenzocyclooctyne (DBCO) endgroups (**Extended Fig. 3a**). Crosslinking of the PEG macromers proceeds via a strain promoted alkyne-azide cycloaddition (SPAAC) bio-click reaction between the azide and DBCO groups, resulting in hydrogels with a final equilibrated shear storage modulus (G’) of ∼0.5 kPa (**Extended Fig. 3a, b**). Intestinal colonies, grown from single ISCs (as in **Extended Fig. 1a**), were encapsulated within this hydrogel formulation modified with the fibronectin-derived Arg-Gly-Asp (RGD) peptide (0.8 mM) and laminin (0.14 mg mL^-1^) (**Fig. 3a, Extended Fig. 3a**) such that there were stoichiometrically equivalent amounts of DBCO and azide in the final formulation.

After hydrogel formation, the nitrobenzyl containing crosslinks were cleaved using a 405 nm confocal laser (60% laser power, 12.64 µs pixel dwell time), softening the matrix at predefined regions adjacent to the organoid and directing crypt budding at specified points in space and time (**Fig. 3a**, **b**, **Extended Fig. 3a, c, d**). In agreement with previous work^15,18^, symmetry breaking and initiation of crypt buds emerged from the spherical organoids in the softened regions, and upon switching the medium to differentiation conditions, mature crypts developed with differentiated Paneth cells (**Fig. 3b**). Similar to *in vitro* organoids grown in Matrigel (**Fig. 1c-i**), organoids in photo-patterned PEG hydrogels also had Paneth cell nuclei with higher lamin A levels, smaller aspect ratios, and closer positioning and alignment with the epithelial base (**Fig. 3c-f**). However, notably, the PEG hydrogels allowed spatiotemporal control of the epithelial forces and buckling, which led to uniform and predictable crypt structures.

We used the photo-patterning method to synchronize the initiation of bud formation and quantify when higher forces upon the nuclei arise after photo-softening of the adjacent matrix. Within just 3 hours after photo-patterning, regions of the epithelia deformed, and protruding buds emerged within the softened regions of the hydrogel matrix. We postulated that at early time points of crypt initiation (≤ 3 hours), changes in lamin A level would be primarily a result of biophysical adaptation of the nuclear mechanics. Indeed, the lamin A to lamin B expression levels increased in the nuclei at the base of the buds, (i.e., at the highly curved regions of the epithelium) (**Fig. 3g**), which contrasted with the cell nuclei interspersed between the buds (i.e., villi regions) (**Fig. 3g, h**). When compared to spherical organoid controls that were not patterned, the lamin A to lamin B ratios were significantly elevated (**Fig. 3g, h**) in the patterned organoids. Together these experiments further reveal that increased epithelial tension within the buds can arise due to local matrix mechanical changes, altering the steady state organoid spherical shape. This then leads to increased nuclear forces and spatially patterned lamin A upregulation, which can then potentiate nuclear shuttling of mechanosensory proteins and downstream signaling necessary for ISC differentiation.

**Figure 3:**
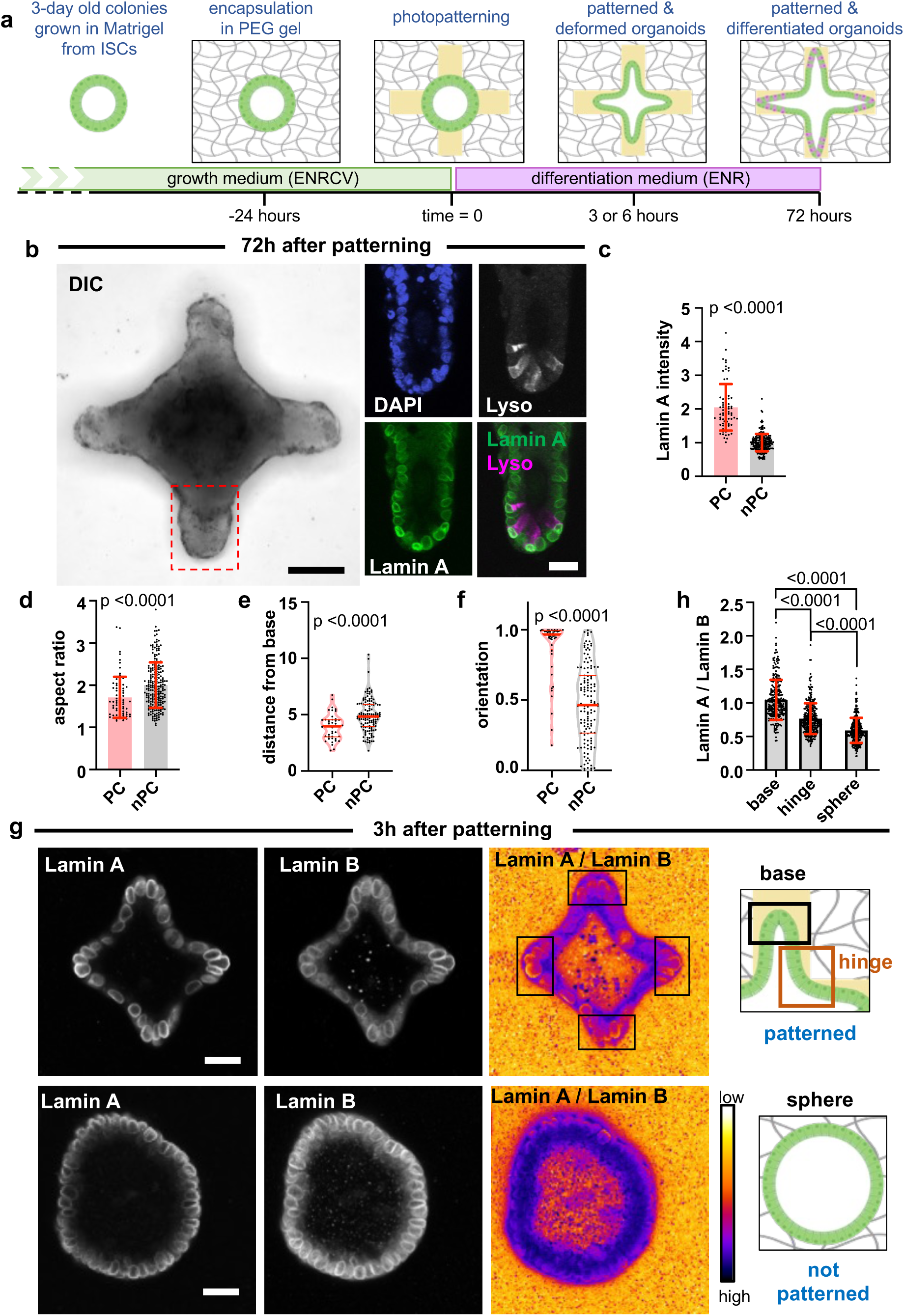
Photo-patternable hydrogels allow spatiotemporal changes in matrix mechanical signaling and reveal increased lamin A over B ratios at regions of high curvature within 3 hours. **a**, Schematic of organoid encapsulation and photo-patterning in PEG based hydrogel. **b,** Representative confocal images of an organoid showing fully developed crypt-like buds 3 days after photo-patterning and incubation in ENR medium. DIC image is a minimum intensity projection while fluorescence images are confocal sections of the region indicated with red box. Images are representative of 3 experiments and scale bars = 50 µm (black) or 20 µm (white). **c**, Lamin A intensities of NE in Paneth (PC) and non-Paneth cells (nPC) within photo-patterned buds. Values normalized to average of nPC intensities in each image. Data are mean ± s.d. from 3 experiments. **d**, Aspect ratio of nuclei in Paneth cells and non-Paneth cells in patterned organoids. Data are mean ± s.d. from 3 experiments. **e**, **f,** Distance of nuclei from epithelium base (e) and orientation of nuclei (f) in patterned organoids from 3 experiments. The red lines indicate the median (thick) and quartiles (thin). **g**, Immuno-stained confocal images of organoids 3 hours after patterning and incubation in ENR medium. Heatmaps show pixel-wise ratio of lamin A over lamin B immunofluorescence. Images are representative of 3 experiments and scale bars = 20 µm. Schematics show the distal region of photo-patterned crypts indicated with black boxes (base) or regions interspersed between the base regions (hinge). Sphere indicates an organoid which is not patterned. **h**, Ratio of lamin A over lamin B immunofluorescence intensities in nuclear lamina of cells occupying base and hinge region of patterned organoid, and those within a spherical organoid. Data are mean ± s.d. from 3 experiments. Statistical analysis was performed using unpaired t-tests after log-transformation (in **c** and **e**), one-way ANOVA followed by Tukey’s multiple comparisons after log-transformation (in **h**), and Mann-Whitney U-test (in **d** and **f**).

### Changes in nuclear envelope precedes canonical YAP mediated mechanotransduction

To identify the possible downstream effects of elevated forces on the nuclei, we probed the time resolved effect of epithelial deformation on activation of YAP. Translocation of the mechanotransducer YAP between the cytoplasm and nuclei is known to be dynamically regulated during intestinal stem cell symmetry breaking events ^26,27^. After binding with transcription factors, nuclear YAP can promote expression of genes like DLL1 in ISCs, which eventually leads to symmetry breaking and generation of Paneth cells^27^. In addition to biochemical regulations, YAP can enter the nuclei after lamin A upregulation^28^ and upon opening of nuclear pores due to increased tension upon the nuclei^9^. Both micropatterned epithelial monolayers^16^ and photo-patterned organoids^17^ have been shown to spatially regulate YAP translocation, verifying its mechanoresponsive nature in the intestinal epithelium. To elucidate the influence of nuclear forces on YAP translocation, we probed the correlation between lamin A levels and nuclear YAP at two different timepoints after photo-patterning (**Fig. 4a**). As observed previously (**Fig. 3g-i**), 3 hours after photo-patterning the levels of lamin A were higher at the base of the epithelial buds (**Fig. 4a**); however, the nuclear to cytoplasmic ratios of YAP were heterogeneous and did not correlate with the lamin A levels at this early timepoint (**Fig. 4a, b**). In contrast, by 6 hours, the YAP localization within the nuclei positively correlated with lamin A levels (**Fig. 4a, c**), suggesting that the nuclei senses forces from epithelial deformation before YAP translocate to the nuclei (**Fig. 4d**).

**Figure 4:**
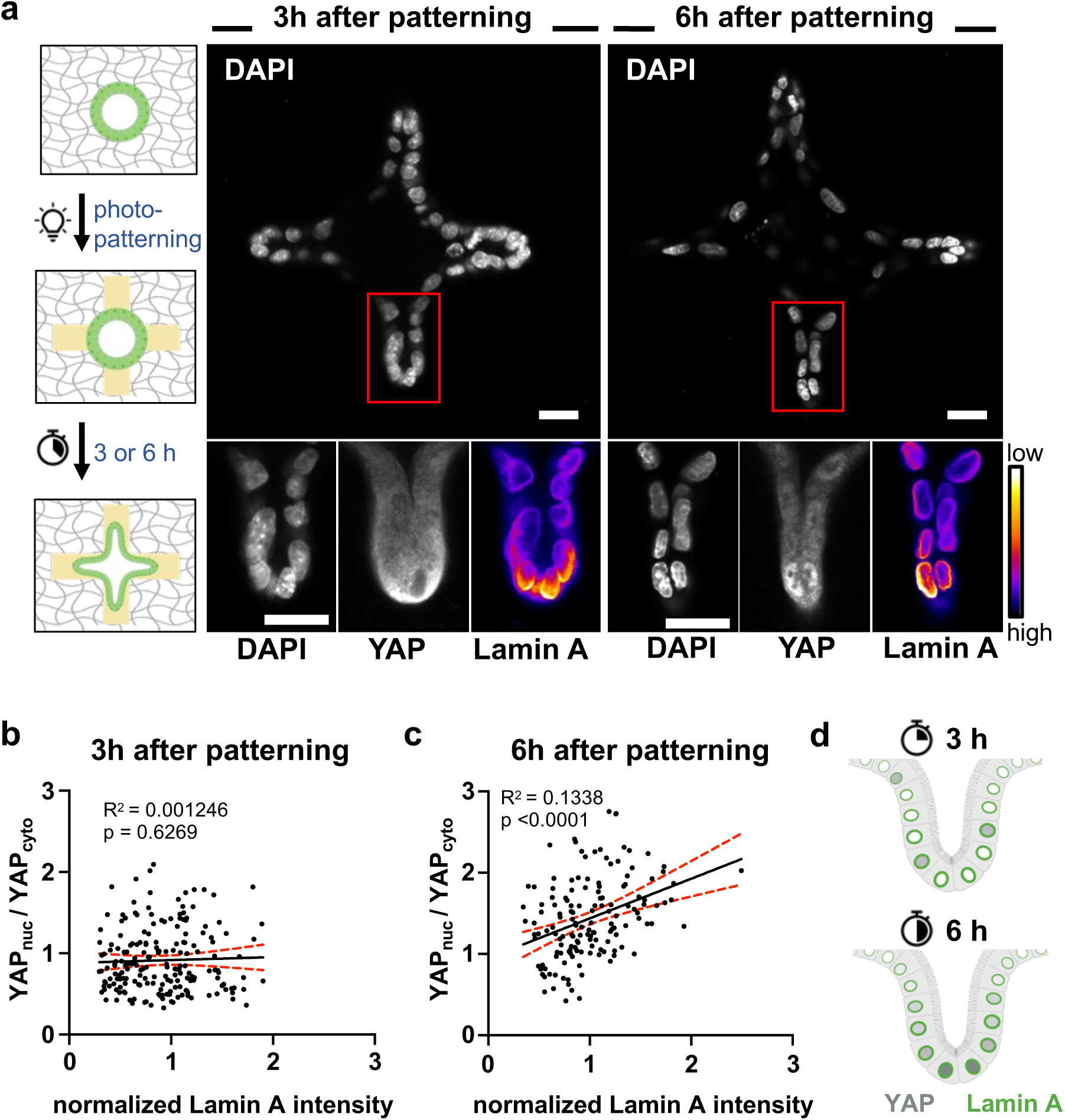
Changes in nuclear lamina precedes YAP translocation. **a**, Schematic shows photo-patterning and immunostaining strategy for YAP and Lamin A correlation analysis at 3h and 6h timepoints. Immuno-stained confocal images of organoids after indicated time of patterning and incubation in ENR medium. Heatmaps show intensity of lamin A immunofluorescence. Images are representative of 3 experiments. Scale bars = 20 µm. **b**, **c**, Ratio of nuclear over cytoplasmic intensity of YAP immunofluorescence plotted against the lamin A intensity in the NE of the corresponding cell 3 hours (b) and 6 hours (c) after photo-patterning. Only cells within the buds were analyzed. Solid lines indicate linear regression and dashed red lines define the 95% confidence intervals from 3 experiments. **d**, Schematic illustration summarizing the sequence of lamin A upregulation and YAP nuclear entry within cells occupying the crypt-like buds in photo-patterned organoids.

### Increased nuclear mechanotransduction alone can promote ISC differentiation

We next asked if epithelial curvature-induced increases in nuclear mechanotransduction would promote ISC differentiation, independent of exogenously added biochemical cues. For this experiment, photo-patterned organoids were maintained in either stem promoting growth medium (ENRCV) or differentiation medium (ENR) and markers for the differentiation of Paneth cells in the crypts were tracked. Irrespective of the medium conditions, organoids demonstrated similar levels of lamin A over lamin B intensities at the base of the buds within 3 hours after photo-patterning (**Extended Fig. 4a, Fig. 5a**). When photo-patterned organoids were maintained in culture for up to 72 hours, we noted differentiated Paneth cells (lysozyme^+^) in both ENR and ENRCV media conditions (**Fig. 5b, c**), and these findings were further verified using *Defa4* and *Dll4* expressing reporter organoid lines (**Extended Fig. 4b, c**). Interestingly, ISCs in unpatterned organoids with spherical geometries displayed spontaneous differentiation in the ENR condition, but rarely produced lysozyme-positive Paneth cells in the ENRCV condition (**Fig. 5b, c**).

We next investigated if increasing nuclear mechanotransduction, by increasing lamin A in the nuclear lamina, would stimulate ISC differentiation in organoids grown in Matrigel and stem promoting conditions. Prior studies had indicated that the retinoic acid inhibitor, AGN-193109 (denoted AGN), increases lamin A expression in a variety of cell types ^8^, and we verified its efficacy in increasing lamin A levels in ISCs with immunostaining (**Extended Fig. 4d**). Indeed, treatment with AGN led to increased number of differentiated Paneth cells and promoted bud formation in ENRCV conditions compared to vehicle control (**Fig. 5d – f**, **Extended Fig. 4e**). Taken together, our data demonstrate that nuclear mechanotransduction alone can regulate stem cell fate, and an upregulation of lamin A levels, either by physical or pharmacological means, can promote differentiation of ISCs to secretory progenitor and Paneth cells, even in stem promoting biochemical conditions.

**Figure 5:**
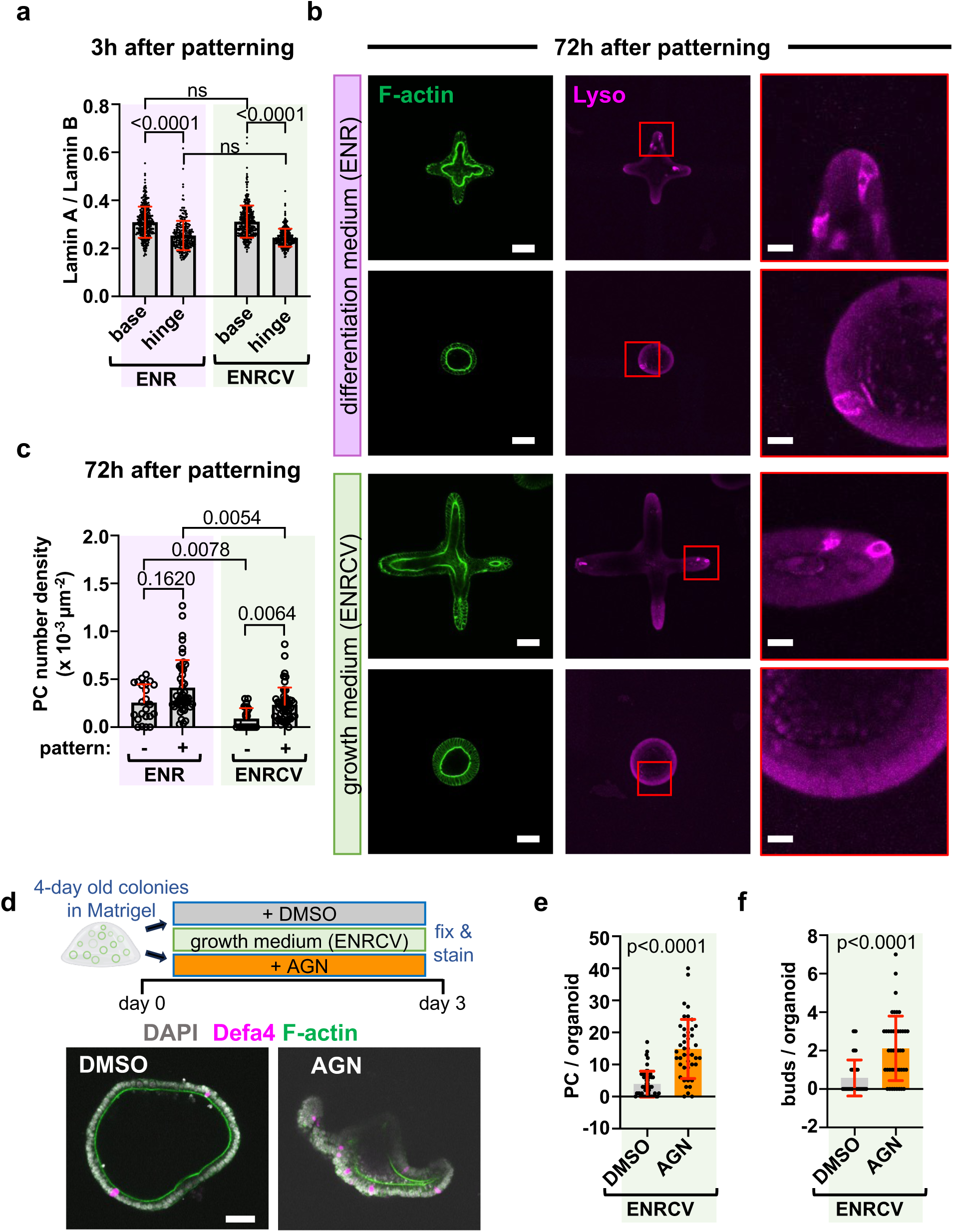
Elevated nuclear mechanotransduction promotes ISC differentiation. **a**, Ratio of lamin A over lamin B immunofluorescence intensities in nuclear lamina of cells after 3h from photo-patterning and incubation in the indicated medium. Base represent cells occupying the distal regions of the photo-patterned crypts and hinge represent cells located in the regions interspersed between the base regions. Data are mean ± s.d. from 3 experiments. **b**, Confocal images of photo-patterned organoids after 3 days of treatment with the indicated medium. Top rows represent patterned organoids while bottom ones were not patterned. Insets show lysozyme-stained Paneth cells from regions marked with red boxes. Images are representative of 3 experiments. Scale bar = 50 µm and 10 µm (in insets). **c**, Number of Paneth cells (PC) in 1000 µm^2^ of epithelial area in patterned (+) or non-patterned, spherical (-) organoids. Data are mean ± s.d. from 3 experiments. **d**, *Top*, schematic of treatment with AGN for organoids grown in Matrigel. *Bottom*, confocal images of *Defa4^Cre^*/*Rosa26^tdTomato^* organoids after 3 days of incubation with stem promoting ENRCV medium and AGN or DMSO (vehicle control). Images are representative of 3 experiments and scale bar = 50 µm. **e**, **f**, Total number of Paneth cells (e) and number of crypt-like buds (f) in organoids treated with AGN or vehicle control in (d). Data are mean ± s.d. from 3 experiments. Statistical analysis was performed using Kruskal Wallis test followed by Dunn’s test (in **a** and **c**) and Mann-Whitney U-test (in **e** and **f**).

### Lamin A is elevated in differentiated cells within mouse intestinal tissues and upon epithelium buckling in human derived organoids

As an *in vivo* comparison, we next examined the NE composition and shape in mouse intestinal crypts to probe the mechanical status of the epithelial nuclei. In sections derived from Defa4^Cre^/Rosa26^tdTomato^ mice, we observed higher lamin A levels in tdTomato^+^ Paneth cells compared to non-Paneth cells residing at the crypt (**Fig. 6a, b**). We noted similar enrichment of lamin A in lysozyme^+^ Paneth cells of intestinal tissues explanted from wild-type mice, highlighting that our observation is not due to Defa4 or tdTomato expression (**Extended Fig. 5a, b**). In line with our *in vitro* observations, lamin B levels in tdTomato^+^ Paneth cells were lower compared to non-Paneth cells (**Fig. 6c**). Lamin B staining of the nuclear lamina also revealed higher nuclear wrinkling in Paneth cells as evidenced by the invagination of nuclear lamina inside the nuclei (**Fig. 6a**) verifying our *in vitro* observations. We then explored if the nuclear response to 3D epithelial deformation was significant and translated to other 3D epithelial models beyond murine crypts. We investigated tumor organoids derived from human pancreatic ductal adenocarcinoma (PDAC) xenografts (**Fig. 6d, e**) and human derived epithelial organoids (HDEs with no mesenchyme) from human intestinal organoids (HIOs) generated from induced pluripotent stem cells (iPSC) (**Fig. 6f, g**). The human PDACs and HDEs were encapsulated in our hydrogel matrices that were then photo-patterned (as in **Fig. 3a**) to alter local forces and initiate epithelium deformations. Within 3 hours after post-patterning, we observed spatial differences in the lamin A to lamin B ratios, elevated at the base of the buds and similar to what we observed in murine ISC organoids (**Fig. 3g, h**). Overall, these results confirm that epithelia of different origins display active changes in nuclear mechanosensing in response to tissue deformation. Mouse tissue explants show that these elevated mechanoresponses are maintained in differentiated Paneth cells, further implicating their role in cell fate specification.

**Figure 6:**
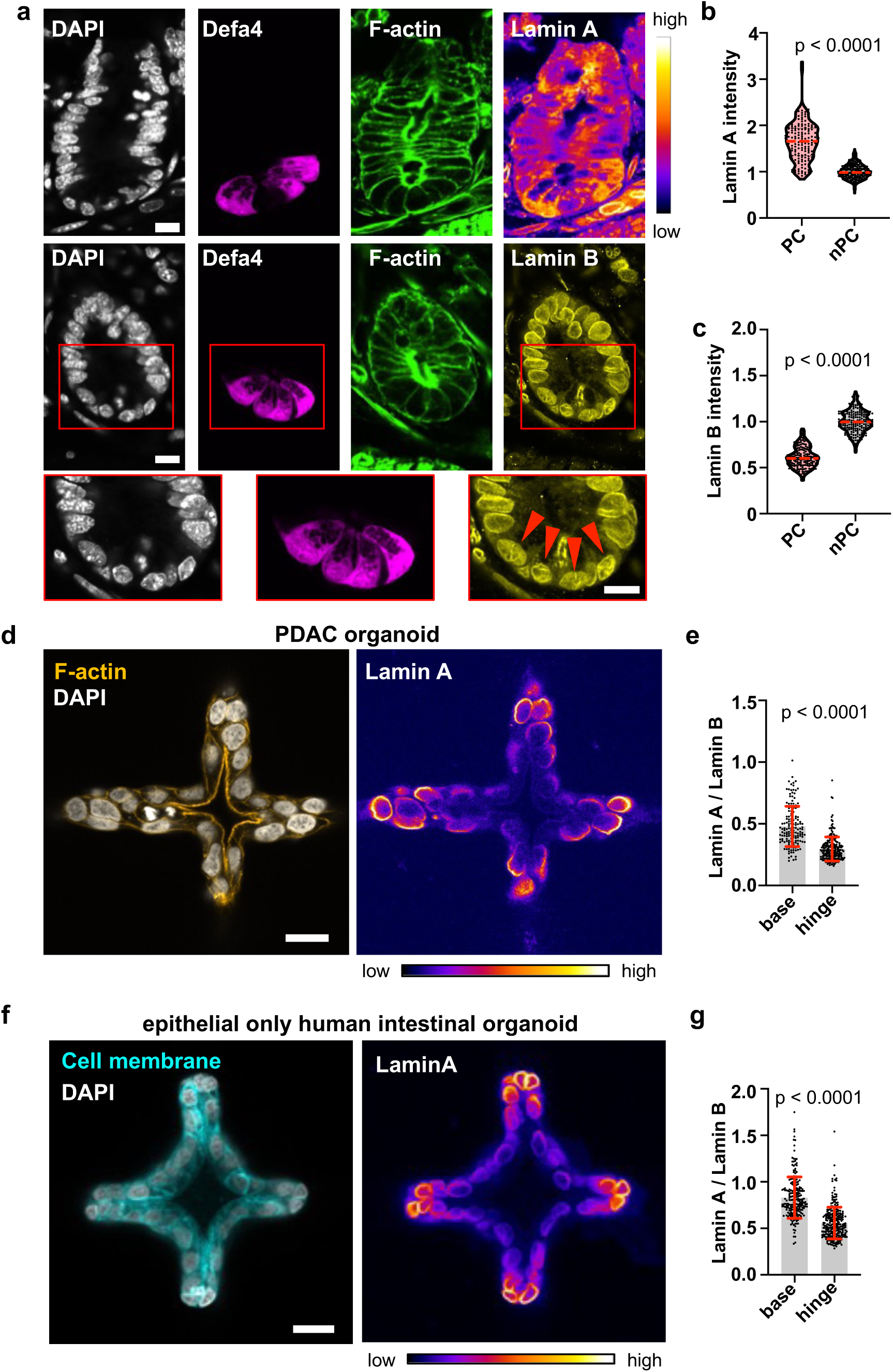
Higher lamin A exists in Paneth cells in mouse intestine and deformed human epithelia. **a**, Confocal images of Defa4^Cre^/Rosa26^tdTomato^ mouse intestinal sections stained with DAPI and lamin A or lamin B. Insets show zoomed in view of the regions indicated by the red boxes. Red arrowheads indicate invaginations of lamina inside Paneth cell nuclei. Scale bars = 10 µm. **b**, **c**, Intensities of lamin A (b) and lamin B (c) in Paneth and non-Paneth cell nuclei within tissue sections. Values normalized to average of nPC intensities in each image. Data are from 3 mice; the dashed red lines indicate the median and dotted black lines indicate the quartiles. **d**, Immuno-stained confocal images of human pancreatic ductal adenocarcinoma (PDAC) derived organoids 3 hours after photo-patterning. Scale bar = 20 µm. **e**, Ratio of lamin A over lamin B immunofluorescence intensities in nuclear lamina of cells occupying the distal regions of the photo-patterned regions (base) or regions interspersed between the base regions (hinge). Data are mean ± s.d. from 3 experiments. **f**, Immuno-stained confocal images of epithelial only human intestinal organoids originating from iPSCs, 3 hours after photo-patterning. Organoids constitutively expressed cell membrane localized tdTomato. Scale bar = 20 µm. **g**, Ratio of lamin A over lamin B immunofluorescence intensities in nuclear lamina of cells occupying the end of the photo-patterned regions (base) or regions interspersed between the patterned regions (hinge). Data are mean ± s.d. from 3 experiments. Images are representative of 3 mice (in a) or 3 experiments (in d and f) and heatmaps quantify the spatial variations in the immunofluorescence intensity of lamin A. Statistical analysis was performed using Mann-Whitney U-tests (in **a**, **b**, and **c**) and unpaired t-test (in **e**).

## Discussion

Nuclear mechanotransduction is increasingly recognized as a key regulator of cell behavior, influencing processes such as proliferation, confined migration, proprioception, and disease progression^29–34^. As a notable example, transient nuclear deformation of reprogrammable fibroblasts by squeezing through microfluidic pores has been shown to enhance cellular differentiation by reducing H3K9me3 and DNA methylation marker 5-methylcytosine, thus releasing epigenetic suppression of cellular reprogramming^35^. Despite such insights at the single-cell level, less is known about nuclear mechanosensing in cells residing in epithelial tissues, where cell-cell contacts are important and changes in the epithelia shape propagate forces throughout the 3D tissue layer. How cells - and the nuclei - interpret these forces is important for proper maintenance of homeostasis in 3D epithelial tissues, and differs fundamentally from isolated mesenchymal cells or those embedded as single cells in 3D matrices

Here we demonstrate that nuclei act as direct sensors of extracellular forces and can determine cell fate using a 3D intestinal epithelium model. We show that stem and differentiated cells, existing in the same niche, can display different nuclear properties and shape, both *in vivo* and *in vitro*. Leveraging the unique properties of photo-responsive matrices, we temporally controlled *de novo* crypt initiation and revealed that forces on the nuclei from early epithelial shape changes manifest in altered NE composition, which is maintained in terminally differentiated Paneth cells. We also demonstrated that increasing force on the nuclei either by photo-patterning or by increasing nuclear mechanotransduction by pharmacological means, induces ISC differentiation in otherwise stem promoting environment. These results reveal the possibility of materials-directed mechanical priming of ISCs towards differentiated lineages. Further, the findings demonstrate the ability to deterministically engineer organoid composition and architecture, potentially increasing their utility for screening of drugs or transplantation for regeneration, where uncontrolled organoid heterogeneity remains a challenge^36^.

The differentiation and survival of ISCs are regulated by the Notch signaling pathway. It is particularly important in directing cells toward absorptive versus secretory lineages through Notch-mediated lateral inhibition and Notch-dependent transcription factor Hes1^14,37,38^. While previous studies showed inhibition of retinoic-X-receptors (RXR) results in undifferentiated regenerative state in organoids^39,40^, villin-targeted deletion of retinoic acid receptor alpha (RAR-α) in the mouse small intestine increases the number of secretory Paneth cells without changing Hes1 activity^39^. Along these lines, inhibition of RAR using the RAR specific inhibitor AGN 193109 in organoids grown under differentiation medium conditions increases the number of secretory and Paneth cells^40^. Here, our results showed that treatment of organoids with AGN, in otherwise stem promoting medium, resulted in higher Paneth cell differentiation and crypt-like bud formation by upregulating lamin A. Thus, our findings suggest an alternate pathway to control the differentiation of ISCs through nuclear mechanotransduction, whose interplay with biochemical cues can be explored in future work.

Further, super resolution 3D imaging utilizing PhotoExM allowed us to uncover and characterize considerable deformations or wrinkles in the NE of Paneth cells both *in vivo* and *in vitro*. Such deformations in 2D cells are typically associated with a relaxed NE due to excess surface area of the nuclei relative to a sphere of equal volume^21,41^. However, single cells embedded in 3D matrices demonstrate increased nuclear wrinkling with increased nuclear YAP^22^. Additionally, transient (< 10 millisecond) squeezing of nuclei in cells passing through microchannels increases their wrinkling^35^, hinting at a context-dependent manifestation of forces on the NE in 2D vs. 3D settings. Consistent with these findings, our laser ablation data confirmed that relaxation of cytoplasmic compressive forces on the Paneth cell nuclei results in a monotonic increase in the nuclear cross-sectional area, further validating that nuclear wrinkling in 3D tissues can imply a compressed nucleus.

The relevance of the nuclear mechanotransduction pathway extends well beyond a murine intestinal organoid model. We confirmed that elevated nuclear lamin A and wrinkling in Paneth cells exists in murine intestinal crypts and also observed mechano-adaptive responses in two other epithelial-based organoids: 1. iPSC derived human intestinal organoids (HIOs) and 2. human PDAC-derived xenograft organoids. *De novo* buckling of both the human organoid epithelia rapidly upregulated lamin A at regions of high tissue curvature and mechanical stress, underscoring a conserved nuclear lamina mediated mechano-adaptation pathway in epithelial nuclei. Together, these findings highlight how nuclear mechanotransduction can potentially regulate cell behavior under varying forces on the epithelium during morphogenesis, healthy organ functioning, and diseases such as fibrosis and cancer.

More broadly, our results position the nucleus as both a mechanical sensor and an active determinant of lineage outcomes in epithelial tissues. Earlier works have shown that uniaxial stretch applied to a flat monolayer of skin epidermal stem/progenitor cells can remodel the heterochromatin to both regulate cell lineage commitment^42^ and protect the genome from stretch induced mechanical damage^43^. Recent studies in the intestinal epithelium also demonstrate the role of H3K36 methylation as an epigenetic regulator of cellular plasticity^44^. Expanding on this, our work raises the possibility that forces imposed on the nucleus during epithelial buckling could directly regulate epigenetic landscapes and dysregulation of the forces might drive aberrant plasticity, dedifferentiation, or neoplastic progression.

## Acknowledgments

We thank Mark Young for help and discussion regarding material synthesis, Michael Blatchley for useful discussion regarding photo-patterning and human iPSC derived organoid culture, Joseph Dragavon for help and advice with confocal microscopy, James D. Orth for help and advice with 2-photon microscopy, and Hannah Larson for help with Olympus Fluoview software. Selected schematics were created with BioRender.com.

We acknowledge funding from the Helen Hay Whitney Foundation (award F1339 to K.B.), the National Institutes of Health (F32-DK142253 awarded to M.Q., T32-HL007822 and F32-HL176073 awarded to A.K., F31-DK126427 awarded to F.M.Y., R01-DK118023 awarded to L.C.S., and R01-DK120921 awarded to K.S.A. and P.J.D.) and the National Science Foundation (Graduate Research Fellowship Program DGE 2040434 awarded to N.R.P., awards 2412520 and 2226157 to T.P.L., and RECODE 2033723 to K.S.A. and P.J.D.). The Organoid and Tissue Modeling Shared Resource (OTMSR) was supported in part by University of Colorado’s Diabetes Research Center (P30-DK116073-01A1), Cancer Center Support Grant (P30 CA046934-34), and the Gates Institute at University of Colorado Anschutz Medical Campus. Selected imaging work was performed at the BioFrontiers Institute’s Advanced Light Microscopy Core (RRID: SCR_018302). The Nikon A1R microscope is supported by NIST-CU Cooperative Agreement award number 70NANB15H226 and the Nikon AXR Laser Scanning Confocal is supported by NIH Grant 1S10OD034320. We acknowledge the Light Microscopy Core Facility, Porter B047, B049, B051, and B059 at the University of Colorado Boulder (RRID:SCR_018993). The opinions, findings and conclusions, or recommendations expressed are those of the authors and do not necessarily reflect the views of any of the funding agencies.

## Author Contributions

K.B. conceptualized and designed the study, performed and analyzed the majority of the experiments, synthesized macromers (with F.M.Y.), and wrote the manuscript. D.L.M., B.E.K., N.R.P., N.P.S., and A.K. performed selected experiments, analyzed data, and edited the manuscript. T.P.L. designed selected experiments (with K.B. and B.E.K.), provided critical input, and edited the manuscript. M.C., M.Q., P.S.M. and L.C.S. generated tissue explants and organoids, performed selected experiments, and edited the manuscript. P.J.D. designed and supervised the study, performed selected experiments, provided critical input, and edited the manuscript. K.S.A. conceptualized, designed and supervised the study, provided critical input, and edited the manuscript.

## Data Availability

The data that support the findings of this study are available within the article and its supplementary information and are available from the corresponding authors upon request. Source data will be provided with this paper during publication.

## Methods

### Animal Studies

All animal procedures were approved by the Institutional Animal Care and Use Committee at the University of Colorado Anschutz Medical Campus (IACUC protocol no. 00084) and the University of Michigan (IACUC protocol no. 00012476). The following mouse strains were used in this study: C57BL/6J (JAX stock no. 000664; The Jackson Laboratory), *Defa4^Cre^* ^20,45^, *Rosa26^tdTomato^*(JAX stock no. 007908; The Jackson Laboratory) ^20,46^, *Dll4-mCherry* BAC transgenic mice^47,48^. Mice were housed in ventilated and automated watering cages with a 12-h light cycle under specific pathogen-free conditions. All experimental animals were adult mice, ≥8 weeks of age and represented both sexes.

### Immunohistochemistry of intestinal sections

Paraffin-embedded tissue sections (6 µm thick) from wild-type control mice were immunostained using a rabbit anti-lysozyme antibody (Invitrogen, PA129680) or rabbit anti-lamin A antibody (Sigma-Aldrich, SAB5700618). Sections were deparaffinized in SafeClear 2 (Fisher Scientific, 23-044192) for 3 minutes at room temperature (RT), then rehydrated through a graded ethanol series (100%, 95%, 70%, 50%, and 0% diluted in deionized water), each for 3 minutes. Antigen retrieval was performed at RT for 15 minutes using a trypsin-based solution (Abcam, ab970). Following permeabilization with 0.1% Triton X-100 in phosphate-buffered saline (PBS), sections were blocked in 5% bovine serum albumin (BSA) dissolved in PBS for 1 hour at RT. Primary antibodies (1:200 dilution in blocking buffer) were applied overnight at 4°C. After washing with 0.05% Tween 20 solution in PBS (PBST) for 10 minutes (4X), sections were incubated with Alexa Fluor 647-conjugated goat anti-rabbit secondary antibodies (1:200; Invitrogen, A21245) for 1 hour at RT. Samples were then washed again with PBST (4X for 10 mins each) and mounted with Fluoro-Gel II containing 4’,6-diamidino-2-phenylindole (DAPI, Electron Microscopy Sciences, 17985-50) and sealed with coverslips.

Tissues embedded in optimal cutting temperature (OCT) compound (Fisher Healthcare, 23-730-571) were sectioned (10 µm thick) using a cryotome (Leica Microsystems) and mounted onto Superfrost Plus Microscope Slides (Fisherbrand). Two PBS washes of 2 minutes were performed followed by 1 hour of permeabilization with 0.2% Triton X-100 in PBS and 1 hour of blocking with 5% BSA and 0.01% Triton X-100 in PBS, both at RT. Primary antibodies raised in rabbit against lamin A (Sigma-Aldrich, SAB5700618) or lamin B (Proteintech, 12987-1-AP) were applied overnight at 4°C (1:200 dilution in blocking buffer). After washing with PBST (4X for 10 mins each), sections were incubated with Alexa Fluor 647-conjugated goat anti-rabbit secondary antibodies (1:200; Invitrogen, A21245), Alexa Fluor 488 conjugated phalloidin (1:400, Invitrogen, A12379) and DAPI (1:400, Sigma-Aldrich, 10236276001) diluted in blocking buffer for 1 hour at RT. Samples were then washed with PBST (4X) and mounted with ProLong Glass Antifade Mountant (Invitrogen, P36984) and sealed with coverslips. Images were acquired with a Nikon AXR laser scanning confocal using a 20X/0.95 NA water immersion objective.

### Generation of mouse organoid lines

Jejunal organoid lines were generated from C57BL/6J mice or mice expressing different fluorescent reporter alleles (*Defa4^Cre/+^/Rosa26^tdTomato/+^; Defa4^Cre/+^/Rosa26^tdTomato/+^/CAG:H2B-GFP* and *Dll4-mCherry*). Briefly, intestinal crypts were isolated from the jejunum (a 2-cm region located 10-12 cm from the gastroduodenal junction). Crypts were plated in ice-cold Reduced Growth Factor Basement Membrane Extract (BME) (Cultrex, R&D Systems) (15 μL) into wells of 48-well plates and grown in 300 μL ENR medium. The basal ENR culture medium (advanced Dulbecco’s modified Eagle medium/F12 [no. 12634020] supplemented with 1X penicillin/ streptomycin [no. 15070063], 10 mM HEPES [no. 15630080], 1X GlutaMAX [no. 35050061], 1X B27 [no. 17504044]; all from Life Technologies), and 1 mM N-acetylcysteine (A7250; Sigma-Aldrich) was supplemented with 50 ng mL^-1^ murine recombinant epidermal growth factor (315-09; PeproTech), 50 ng mL^-1^ murine recombinant Noggin (250-38; PeproTech) or Noggin conditioned medium (20% final volume), and R-spondin 2 conditioned medium (10% final volume). When needed, enteroids were grown in ENRCV medium, in which ENR medium was supplemented with 3µM CHIR99021 (C) and 10mM valproic acid (V). For the first 2 days after passaging, organoid medium was supplemented with 10 µM Y27632. Medium was replaced every 2–3 days.

### Media preparations for organoid experiemnts

Organoid “basal medium” was prepared from Advanced DMEM/F12 (Gibco, 12634010) supplemented with GlutaMAX (Gibco, 35050061), 10 mM HEPES (Gibco, 15630080), and 100 U mL^-1^ and 100 µg mL^-1^ of penicillin and streptomycin, respectively (Gibco, 15140122). Basal medium was supplemented with 5% v/v <u>R</u>-spondin 2-conditioned medium (Organoid and Tissue Modeling Shared Resource, University of Colorado Anschutz Medical Campus), N-2 (Gibco, 17502048), B-27 (Gibco, 17504044), 50 ng mL^-1^ recombinant murine <u>e</u>pidermal growth factor (EGF, PeproTech/Gibco, 315-09-500UG), 100 ng mL^-1^ recombinant murine <u>n</u>oggin (PeproTech/Gibco, 250-38-20UG), 3 µM <u>C</u>HIR 99021 (Tocris Bioscience, 4423), 1 mM <u>V</u>alproic acid sodium salt (VPA, Millipore Sigma, P4543-10G) and 1 mM N-Acetyl-L-cysteine (NAC, Sigma-Aldrich, A9165-5G) to obtain “complete growth medium” or <u>ENRCV</u> medium. Experiments with ENR medium were performed by using complete growth medium without CHIR 99021 and VPA. In selected experiments, the following pharmacological agents were added to the culture medium: 4 µM Cytochalasin D (CytoD, Sigma-Aldrich), 1 µM AGN 193109 (Tocris Bioscience, 5758).

### ISC colony formation and differentiation

Organoids cultured in complete medium were harvested by mechanically disrupting the Matrigel using a pipette tip in cold basal medium, followed by centrifugation at 100 g for 5 minutes to obtain a pellet of organoids. The pellet was enzymatically dissociated into single cells by incubating at 37°C for 10 minutes in 1 mL of TrypLE Express (Life Technologies) supplemented with ∼10 mg of DNase I (Millipore Sigma), 1 mM N-acetyl cysteine (NAC, Sigma-Aldrich), and 10 μM Y-27632 (Stemgent). After incubation, the cell suspension was diluted with 1 mL fetal bovine serum (FBS, Gibco) and 3 mL of basal medium, then passed through a 40 μm cell strainer to remove multicellular aggregates. The resulting single-cell suspension was centrifuged at 200 g for 5 minutes, and the cell pellet was resuspended in Matrigel. Matrigel droplets (10 µL) containing the single cells were dispensed into a pre-warmed 48-well Nuncon Delta surface treated plate and allowed to solidify at 37°C for 20 minutes. Complete medium supplemented with 2.5 μM thiazovivin (Stemgent) was then added to each well (300 µL). Intestinal organoids were allowed to develop from single cells over the next 3 days.

For experiments with differentiated organoids, 3-day old colonies were harvested from Matrigel islands using the process outlined above. The colonies were then resuspended in fresh Matrigel and 5 µL droplets were dispensed into 35 mm glass bottom dishes (Ibidi) or glass bottom plates (24 well, Cellvis). After solidification of the Matrigel, the organoids were grown for another day in complete ENRCV medium. After 24 hours, the medium was replaced with ENR medium to promote differentiation and was changed every 2 days.

### Immunofluorescent staining of organoids

Organoids embedded in Matrigel were first washed with PBS followed by incubation with 4% paraformaldehyde (PFA, Electron Microscopy Sciences) in PBS for 1 hour at 37°C. Organoids embedded in PEG hydrogels were treated with 4% PFA for 20 minutes at RT. Fixed samples were permeabilized with 0.2% Triton X-100 (Sigma-Aldrich) in PBS for 1.5 hours at RT, followed by blocking for 1.5 hours at RT in a solution of 5% goat serum (Gibco, 16210072) and 0.01% Triton X-100 in PBS. Organoids were then incubated overnight at 4°C with primary antibodies diluted 1:200 in the blocking buffer. The following day, samples were washed at 4°C with fresh PBS every hour for 5 hours to remove unbound antibody. Subsequently, organoids were incubated overnight at 4°C with DAPI (1:400) and secondary antibodies (1:200) diluted in blocking buffer. In selected experiments samples were also treated with Alexa Fluor 488 conjugated phalloidin (1:400, Invitrogen, A12379). After secondary staining, samples were washed at 4°C with fresh PBS every hour for 3 hours and stored at 4°C prior to imaging. The primary antibodies used were anti-lysozyme (rabbit, Thermo Fisher Scientific, PA129680), anti-lamin A/C (4C11) (mouse, Cell Signaling, 4777), anti-lamin B (rabbit, Proteintech, 12987-1-AP), anti-YAP (D8H1X) (rabbit, Cell Signaling, 14074) and anti-DLL1 (sheep, Cell Signaling, 4777). Secondary antibodies used were Alexa Fluor 647 goat anti-rabbit (Invitrogen, A21245), Alexa Fluor 488 goat anti-rabbit (Invitrogen, A11008), Alexa Fluor 488 goat anti-mouse (Invitrogen, A11001), Alexa Fluor 647 goat anti-mouse (Invitrogen, A21235) and Alexa Fluor 647 donkey anti-sheep (Invitrogen, A21448).

### Photo-expansion Microscopy

For 3D super resolution imaging of organoids, a protocol developed previously was adapted^23,24^. Organoids from ISC colonies were embedded in 5 µL Matrigel droplets on 12 mm circular cover glasses and were kept in ENRCV medium for 24 hours before switching to differentiation medium (ENR). After 3 days of differentiation, samples were fixed and immunolabeled for lysozyme using the process described in the previous section. Antibodies used were polyclonal rabbit anti-lysozyme EC 3.2.1.17 (Dako, A0099, 1:1000 dilution) as primary and Alexa Fluor Plus 488 goat anti-rabbit (Invitrogen, A48282,1:200 dilution) as secondary. Subsequently, samples were treated with 0.1 mg mL^-1^ Acryloyl-X in PBS for 6 hours at RT and then washed 3 times with PBS for 20 minutes each. A solution was prepared with sodium acrylate (16 wt%), 8-arm, 10 kDa PEG-thiol (6 wt%), PEG-bisacrylamide (0.875 wt%), acrylamide (3 wt%) and lithium phenyl-2,4,6-trimethylbenzoylphosphinate (LAP, 0.2 wt%) in PBS supplemented with 2 M NaCl, to obtain the PhotoExM formulation. The PhotoExM formulation used was optimized for 4.5X expansion prior to adding to samples. Coverslips containing the Matrigel island were transferred into 5.5 cm Petri dishes, excess liquid around the Matrigel droplets was removed, and the islands were incubated with the PhotoExM solution for 30 minutes (2X) by adding the solution on top of the Matrigel. After permeation with the PhotoExM solution, Rain-X-coated glass slides were placed on top of the Matrigel at 1-2 mm separation from the bottom coverslip and the samples were irradiated with 365 nm light (4.5 mW cm^-2^) for 70 seconds. Samples were then digested overnight at 37°C with proteinase K (8 units mL^-1^) dissolved in digestion buffer (100 mL of which was prepared by mixing 0.5 g Triton-X, 27 mg EDTA, 4.67 g NaCl, and 5 mL 1M Tris aqueous solution in DI water). Samples underwent a first round of expansion initiated by a 20-minute wash in DI water, followed by staining with DAPI (1:500) for 1 hour in DI water. Final expansion was achieved through two additional 20-minute DI water washes. The expanded gels were cut to appropriate dimensions using a rectangular cover glass and then positioned on a glass bottom plate (6 well, Cellvis) for imaging.

### Live imaging of organoids

Live tracking of organoid nuclei was performed using a Nikon laser scanning confocal microscope fitted with an Oko Labs environmental chamber. *Defa4^Cre^*/*Rosa26^tdTomato^* Organoids, differentiated in Matrigel for 3 days, were incubated with Hoechst 33342 (NucBlue Live ReadyProbes Reagent, Invitrogen, 1:200) at 37°C for 30 minutes before imaging with a 20X/0.95 NA water immersion objective. The organoids were maintained at 37°C and 5% CO_2_ while imaging with a 405 nm laser. Multiple positions were imaged in time-lapse sequences using the Nikon Perfect Focus System (PFS) and Nikon Elements imaging software.

### Laser ablation of organoids

Laser ablation and imaging of cells in organoids were performed with an Olympus FVMPE-RS Twin Laser Multi-Photon microscope connected to two tunable, pulsed IR lasers: a Spectra Physics Insight X3 (680-1300 nm) and Spectra Physics MaiTai eHP DeepSee (690-1040 nm). *Defa4^Cre^*/*Rosa26^tdTomato^*organoids with or without a H2B-GFP reporter were differentiated in Matrigel for 3 days before imaging using a 25X/1.05 NA water immersion objective. For organoids without H2B-GFP, the nuclei were stained using Hoechst 33342 following the process reported above. Imaging and ablation sequences were controlled by the Olympus Fluoview software and lasers were tuned at 1100 nm (for tdTomato), 900 nm (for GFP), and 800 nm (for Hoechst). The MaiTai laser system, operating at 2 W and a repetition rate of 80 MHz, was tuned to 800 nm with a 2 µs pixel^-1^ scan speed to ablate a region of interest at the apical surface of the cell using a linear stimulation ROI drawn across the membrane. After stimulation, the nuclei were imaged every 30 seconds for a total of 5 minutes.

### Synthesis of DBCO-functionalized 40 kDa PEG

Eight-arm PEG macromers bearing DBCO groups (PEG-8-DBCO) were prepared via amide coupling between DBCO-C6 acid (100 mg, 0.3 mmol; Click Chemistry Tools) and 8-arm PEG-amine (0.5 g, 0.025 mmol; MW 40,000; JenKem USA). The reaction was facilitated by the coupling agent (1-[bis(dimethylamino)methylene]-1H-1,2,3-triazolo[4,5-b]pyridinium 3-oxid hexafluorophosphate (HATU, 121 mg, 0.32 mmol; ChemPep) in the presence of N,N-diisopropylethylamine (0.175 mL, 1 mmol; Sigma Aldrich) as the base. The mixture was purged with argon and allowed to react overnight under inert conditions. Following reaction completion, the product was purified by repeated precipitation in chilled diethyl ether (3X), and the resulting solid was collected by centrifugation. The crude product was then dissolved in deionized water and subjected to dialysis (8 kDa molecular weight cutoff; Spectrum Chemical) for 48 hours against DI water. The final product was obtained after lyophilization.

### Synthesis of nitrobenzyl azide-functionalized 40 kDa PEG

Nitrobenzyl azide-functionalized 8-arm PEG macromers were synthesized through a multi-step modification of 8-arm PEG-amine (1 g, 0.025 mmol; MW 40,000; JenKem USA). In the first step, a photolabile Fmoc-protected nitrobenzyl linker (218 mg, 0.42 mmol; Advanced ChemTech) was conjugated to PEG-amine using HATU (152 mg, 0.4 mmol; ChemPep) as the coupling agent and 4-methyl morpholine (107 mg, 0.8 mmol; Sigma Aldrich) as the base. The reaction was conducted under an argon atmosphere and allowed to proceed overnight. The crude product was purified by precipitation in ice-cold diethyl ether, collected by centrifugation, and redissolved in a solution of 80% N,N-dimethylformamide (DMF) and 20% piperidine for Fmoc deprotection.

After 24 hours of deprotection, the resulting intermediate product was precipitated again in ice-cold diethyl ether and collected as a solid. This intermediate was then reacted with 4-azido butanoic acid (62 mg, 0.48 mmol), prepared according to previously published protocols^49^, using the same coupling conditions (HATU and 4-methylmorpholine under argon overnight in DMF). The final PEG-8-NBA macromer was precipitated in cold diethyl ether, dissolved in minimal DI water, dialyzed for 48 hours (8 kDa MW cut off), and lyophilized prior to subsequent use.

### Characterization of hydrogel mechanical properties and degradation

Photo-degradable hydrogels containing nitrobenzyl ether linkages were formed via SPAAC by combining stock solutions of 10 wt% PEG-8-DBCO and 20 wt% PEG-8-NBA in phosphate-buffered saline (PBS) at 4°C. The solutions were mixed to yield a final polymer concentration of 2.5 wt% PEG in PBS, containing 2.9 mM DBCO groups and 2.1 mM azide groups. The precursor solution was vortexed for 3 seconds and 25 µL drops were immediately added between two glass slides treated with Sigmacote (Sigma-Aldrich) and separated by 480 µm silicone spacers. The gels were allowed to polymerize for 10 minutes before they were transferred to a well plate with PBS (1 mL). After reaching equilibrium overnight in PBS, the gels were transferred to a rheometer (Ares-G2, TA Instruments) configured with a 405 nm laser. Initial shear storage (G’) and loss (G”) moduli were measured by operating the instrument at 1% strain and 1 Hz frequency before turning on the laser (50 mW cm^-2^) to track the mechanical properties of the gel as the network degraded.

To further characterize the degradation during confocal laser scanning, rectangular gels of 250 µm thickness were formed on 35 mm glass bottom dishes (Ibidi) and 40 µm wide X 450 µm long regions at the edge of the gels were irradiated with a Zeiss LSM-710 microscope with the 405 nm laser operating at different power levels between 5% and 100%. The scanning speed was fixed at 12.64 μs pixel dwell time. After patterning, the gels were merged in 1 mg mL^-1^ 2 mega dalton FITC-Dextran (Sigma-Aldrich) solution in PBS. Images of the perfused channels were acquired using a Nikon AXR laser scanning confocal microscope Nikon Spatial Array Confocal (NSPARC) detector with a 10X/0.45 NA air objective.

### Encapsulation of organoids in PEG hydrogels

Hydrogel precursor solutions were made by first diluting DBCO-functionalized PEG in basal medium. Azide-functionalized Arg-Gly-Asp (RGD) peptide (0.8 mM) was added and allowed to react for 5 minutes at room temperature, then the solution was chilled on ice. Laminin (0.14 mg mL^-1^; Corning) was subsequently added and mixed thoroughly. Three-day-old organoid colonies were recovered from Matrigel using cold basal medium and mechanical agitation, followed by centrifugation at 100 g for 5 minutes. The resulting organoid pellet was resuspended in cold basal medium and gently mixed with the chilled hydrogel precursor solution. Hydrogel polymerization was initiated by adding azide nitrobenzyl ether-functionalized PEG to achieve a final polymer concentration of 2.5 wt%. The mixture was vortexed for 3 seconds and immediately dispensed as 6 μL droplets between Sigmacote-treated glass slides and thiol-functionalized coverslips, separated by 250 μm spacers. After allowing 10 minutes for gelation at room temperature, the coverslips with the adhered hydrogels were transferred into a 48-well plate and submerged in 300 μL of complete medium.

### Photo-patterning of encapsulated organoids

Individual gels adhered to coverslips were removed from culture medium one day after organoid encapsulation and placed upside down on a circular rubber stand for imaging and photo-patterning. Photo-patterning was performed using a Zeiss LSM 710 upright laser scanning confocal microscope. Four regions of interest (ROIs), positioned symmetrically around each organoid colony, were scanned within a single optical plane using a 405 nm laser, set at 60% laser power (2.72 mW) with a pixel dwell time of 12.64 μs and a pixel resolution of 420 nm. Approximately 30 colonies per gel were patterned in this manner. After patterning, gels were transferred into fresh medium (ENR or ENRCV) and incubated at 37°C with 5% CO₂.

### Human PDAC tissue and patient-derived xenografts

Patient derived tumor samples were collected from consenting PDAC patients undergoing surgical resection or tumor biopsy at the University of Colorado Cancer Center with approval by the Colorado Multiple Institutional Review Board (08–0439). Tissue samples were confirmed to be tumor or normal based on pathologist assessment.

All animal work was approved by the Institutional Animal Care and Use Committee at the University of Colorado Anschutz Medical Campus. Fresh PDAC tissue samples were used to generate patient-derived xenograft (PDX) models as described previously^50^.

### Human PDAC organoid line

The PDAC269 organoid line was generated from an established patient-derived xenograft (PDX) PDAC tumor model as previously described with some modifications^51–53^. Briefly, PDAC tumor tissue was washed vigorously and minced with surgical scissors. The fragments were then digested with digestion medium [Advanced DMEM/F12 (Gibco) containing 1% fetal bovine serum (FBS), 10mM HEPES (Gibco), 2mM GlutaMAX (Gibco) and 1% Penicillin/Streptomycin (Gibco) with 1.5mg mL^-1^ collagenase (Sigma, C2139), 125µg mL^-1^ dispase, 0.1mg mL^-1^ DNAse1 (EMD Millipore) and 10μM Y27632] at 37°C for 30 min. The digested cells were washed with wash medium (DMEM containing 10% FBS, 10mM HEPES, 2mM GlutaMAX, 1% penicillin/streptomycin and 10μM Y27632) to inactivate the digestive enzymes and then filtered through a 100μm strainer to remove large undigested fragments prior to culture. Isolated pancreatic cells were embedded in Basement Membrane Extract (BME, Biotechne 9mg mL^-1^) and medium was refreshed every 2–3 days. Basal culture medium contained advanced DMEM/F12 supplemented with 1% penicillin/streptomycin, 10mM HEPES, 2 mM GlutaMAX, 1X B27 (Invitrogen), 10 nM gastrin, 1mM N-acetylcysteine and 10mM nicotinamide. Human Pancreatic Stem Cell (HPSC) medium contained basal culture medium supplemented with 50 ng mL^-1^ mouse recombinant EGF, 50% Wnt3A, Noggin, R-Spondin-3 (WNR) conditioned medium, 500 nM A83-01 (Tocris), 10 μM SB202190 (Sigma) and 100 mg mL^-1^ Primocin (Vitrogen). Organoid medium was refreshed every two days and organoids were passaged as needed based on organoid growth and confluency. Frozen stocks of all PDAC organoid lines were prepared and all experiments were performed within a 10–20 passage window. Organoid cultures were routinely tested for mycoplasma.

### iPSC culture and intestinal organoid differentiation

Human intestinal organoids (HIOs) were differentiated, maintained, and passaged following established protocols^54,55^. Induced pluripotent stem cells (iPSCs; line iC4-4 with constitutive membrane tdTomato expression driven by the CAGG promoter and inserted into the AAVS safe-harbor locus; CU Anschutz Organoid and Tissue Modeling Shared Resource) were expanded at 37°C with 5% CO_2_ on six-well plates coated with stem cell–qualified Cultrex (R&D Systems, 3434-005-02). Cells were cultured in mTeSR Plus medium (Stem Cell Technologies, 100-0276) supplemented with penicillin–streptomycin (100 U mL^-1^; Gibco, 15120122). Colonies were clump-passaged using 0.5 mM EDTA in PBS (Invitrogen, 15575020).

Directed differentiation toward definitive endoderm was performed over three days using Activin A (100 ng mL^-1^; Cell Guidance Systems, GFH6-100) in RPMI 1640 (Gibco, 11875199) with penicillin–streptomycin (100 U mL^-1^). HyClone defined FBS (Cytiva, SH3007003) was added at 0%, 0.2%, and 2% on days 1, 2, and 3, respectively. From day 3, endoderm monolayers were exposed to hindgut induction medium consisting of FGF-4 (500 ng mL^-1^; R&D Systems, 235-F4) and CHIR99021 (2 µM; Tocris Bioscience, 44-231-0) in the definitive endoderm medium. Between days 4–6, budding spheroids were harvested from the monolayer, encapsulated in Cultrex BME (R&D Systems, 3533-010-02), and maintained in intestinal growth medium. The intestinal growth medium contained Advanced DMEM/F12 (Gibco, 12634028), B27 supplement (1X; Gibco, 17504044), GlutaMAX (1X; Gibco, 35050061), penicillin–streptomycin (100 U mL^-1^), HEPES (15 mM; Gibco, 15630080), epidermal growth factor (EGF, 100 ng mL^-1^; R&D Systems, 236-EG-200), noggin (100 ng mL^-1^; R&D Systems, 3344-NG-050), and 5% (v/v) R-Spondin2 conditioned medium (CU Anschutz Organoid and Tissue Modeling Shared Resource). Media were refreshed every 3–4 days. HIOs were first passaged at approximately day 14, and by day 28 of 3D outgrowth, they displayed a robust epithelium surrounded by supporting mesenchyme.

HIO epithelial only organoids (HDEs) were established as previously described^56^ with minor modifications. Briefly, to generate HIO epithelial only organoids (HDEs), HIOs were incubated with dispase for 30 minutes. Dispase was removed and replaced by 50% FBS/advanced DMEM/F12 medium followed by mild TrypLE digestion and mechanical disruption to release the mesenchymal layer. TrypLE digestion was repeated until epithelial only organoid structures were observed and collected manually on a phase microscope by pipet. Isolated HDEs were washed, embedded in Cultrex BME, and propagated.

### HIO epithelial only organoid (HDE) culture

HDEs were maintained with HDE growth medium in Cultrex BME. The medium consisted of 50% 2X basal medium (Advanced DMEM/F12 supplemented with B27 [2X; Gibco, 17504044], N-2 [2X; Gibco, 17502048], HEPES [20 mM], GlutaMAX [2X], penicillin–streptomycin [200 U mL^-1^], nicotinamide [20 mM; Sigma, N0636], and N-acetyl-L-cysteine [2 mM; Sigma, A9165]) combined with 50% LWRN conditioned medium (CU Anschutz Organoid and Tissue Modeling Shared Resource). This was further supplemented with A83-01 (2.5 µM; Tocris Bioscience, 293910), CHIR99021 (4 µM), Thiazovivin (2.5 µM; Selleckchem, S1459), SB431542 (100 nM; Tocris Bioscience, 16141), and epidermal growth factor (EGF, 100 ng mL^-1^).

Culture medium was refreshed every 2–3 days, and HDEs were passaged once per week. For passaging, organoids were released from the Cultrex BME, mechanically dissociated, and re-embedded in fresh Cultrex BME.

### Image analysis

ImageJ (National Institutes of Health) was used for the following analyses unless otherwise stated.

<u>Nuclear morphology and NE composition analysis</u>: The nucleus channel was used to manually trace boundaries around the nuclei using the *Freehand selections* tool. Then *Analyze* -> *Measure* function was used to obtain the aspect ratio, cross sectional area, and solidity values of the nuclei. For measurement of lamin A or lamin B intensities, the boundaries were first extended by 0.8 μm using *Edit* -> *Selection* -> *Make Band*… tool to allow the resulting bands to overlap with the NE staining. Then, the mean intensity inside the band was measured on the corresponding channel for lamin A or B. For analysis of DLL1 intensity, the initial boundaries were drawn at the cell periphery prior to band extension and cells with lamin A intensities higher than 12000 arbitrary units were considered.

<u>Nuclear position and orientation analysis</u>: The boundaries of the nuclei were outlined as explained above, converted to masks using the *Create Mask* tool, and then saved as .tif files. The crypt boundaries were then traced on the F-actin (Phalloidin) channel using the *Freehand Line* tool, converted to masks, transformed using the *Skeletonize* tool, and then saved as .tif files. A custom written MATLAB (MathWorks) code was then used to identify: 1) the shortest distance between the centroid of each nucleus and the corresponding crypt boundary; 2) the orientation of the major axis of each nucleus with respect to the tangent at the closest point on the crypt boundary.

<u>Cell number density analysis</u>: Lysozyme or Defa4 positive cells in a 50 μm tall confocal z stack were manually counted to get the total number of cells in the epithelium. Then, a custom written MATLAB (MathWorks) code was used to identify the epithelium boundary at each z stack. This was then used to calculate the total surface area of the measured epithelium. Finally, the total number of cells from each organoid was divided by the surface area of the corresponding epithelium to obtain the cell number density.

### Statistics and reproducibility

Unless otherwise mentioned, data represent the mean ± s.d. of ≥3 experiments (reflecting independent biological replicates) with ≥10 organoids analyzed per condition and data points denote values from each cell. The D’Agostino–Pearson omnibus normality test was used to determine whether data were normally distributed. Datasets with Gaussian distributions were compared using two-tailed Student’s t-tests and one-way ANOVA followed by Tukey’s post hoc test. For log normal distribution, statistical comparison was made after logarithmic transformation of the data, followed by unpaired two-tailed Student’s t-tests or one-way ANOVA followed by Tukey’s post hoc test. For non-Gaussian distributions, nonparametric Mann–Whitney U-tests were used comparing two conditions, and comparisons for more than two groups were performed using Kruskal–Wallis tests followed by Dunn’s multiple-comparison test. Analysis was performed using GraphPad Prism 9 or 10 (GraphPad Software). P < 0.05 was considered to be statistically significant.

## References

1 Guillot, C. & Lecuit, T. Mechanics of epithelial tissue homeostasis and morphogenesis. Science 340, 1185–1189 (2013). 10.1126/science.1235249

2 Nelson, C. M. On Buckling Morphogenesis. J Biomech Eng 138, 021005 (2016). 10.1115/1.4032128

3 Fiore, V. F., Almagro, J. & Fuchs, E. Shaping epithelial tissues by stem cell mechanics in development and cancer. Nat Rev Mol Cell Biol 26, 442–455 (2025). 10.1038/s41580-024-00821-0

4 Armon, S., Bull, M. S., Moriel, A., Aharoni, H. & Prakash, M. Modeling epithelial tissues as active-elastic sheets reproduce contraction pulses and predict rip resistance. Communications Physics 4, 216 (2021). 10.1038/s42005-021-00712-2

5 Vicente, F. N. & Diz-Muñoz, A. Order from chaos: How mechanics shape epithelia and promote self-organization. Current Opinion in Systems Biology 32–33, 100446 (2023). 10.1016/j.coisb.2023.100446

6 Kalukula, Y., Stephens, A. D., Lammerding, J. & Gabriele, S. Mechanics and functional consequences of nuclear deformations. Nat Rev Mol Cell Biol 23, 583–602 (2022). 10.1038/s41580-022-00480-z

7 Uhler, C. & Shivashankar, G. V. Regulation of genome organization and gene expression by nuclear mechanotransduction. Nat Rev Mol Cell Biol 18, 717–727 (2017). 10.1038/nrm.2017.101

8 Swift, J. et al. Nuclear lamin-A scales with tissue stiffness and enhances matrix-directed differentiation. Science 341, 1240104 (2013). 10.1126/science.1240104

9 Elosegui-Artola, A. et al. Force Triggers YAP Nuclear Entry by Regulating Transport across Nuclear Pores. Cell 171, 1397–1410 e1314 (2017). 10.1016/j.cell.2017.10.008

10 Blatchley, M. R. & Anseth, K. S. Middle-out methods for spatiotemporal tissue engineering of organoids. Nat Rev Bioeng 1, 329–345 (2023). 10.1038/s44222-023-00039-3

11 Zhu, G., Hu, J. & Xi, R. The cellular niche for intestinal stem cells: a team effort. Cell Regen 10, 1 (2021). 10.1186/s13619-020-00061-5

12 Ritsma, L. et al. Intestinal crypt homeostasis revealed at single-stem-cell level by in vivo live imaging. Nature 507, 362–365 (2014). 10.1038/nature12972

13 Spit, M., Koo, B. K. & Maurice, M. M. Tales from the crypt: intestinal niche signals in tissue renewal, plasticity and cancer. Open Biol 8 (2018). 10.1098/rsob.180120

14 Baghdadi, M. B. et al. PIEZO-dependent mechanosensing is essential for intestinal stem cell fate decision and maintenance. Science 386, eadj7615 (2024). 10.1126/science.adj7615

15 Yavitt, F. M. et al. Engineered epithelial curvature controls Paneth cell localization in intestinal organoids. Cell Biomater 1 (2025). 10.1016/j.celbio.2025.100046

16 Young, M. W. et al. Synthetic photoresponsive hydrogels enable in situ control over murine intestinal monolayer differentiation and crypt formation. Adv Funct Mater 35 (2025). 10.1002/adfm.202413778

17 Yavitt, F. M. et al. In situ modulation of intestinal organoid epithelial curvature through photoinduced viscoelasticity directs crypt morphogenesis. Sci Adv 9, eadd5668 (2023). 10.1126/sciadv.add5668

18 Gjorevski, N. et al. Tissue geometry drives deterministic organoid patterning. Science 375, eaaw9021 (2022). 10.1126/science.aaw9021

19 Wang, M., Ivanovska, I., Vashisth, M. & Discher, D. E. Nuclear mechanoprotection: From tissue atlases as blueprints to distinctive regulation of nuclear lamins. APL Bioeng 6, 021504 (2022). 10.1063/5.0080392

20 Jones, J. C. et al. Cellular Plasticity of Defa4(Cre)-Expressing Paneth Cells in Response to Notch Activation and Intestinal Injury. Cell Mol Gastroenterol Hepatol 7, 533–554 (2019). 10.1016/j.jcmgh.2018.11.004

21 Dickinson, R. B., Abolghasemzade, S. & Lele, T. P. Rethinking nuclear shaping: insights from the nuclear drop model. Soft Matter 20, 7558–7565 (2024). 10.1039/d4sm00683f

22 Cosgrove, B. D. et al. Nuclear envelope wrinkling predicts mesenchymal progenitor cell mechano-response in 2D and 3D microenvironments. Biomaterials 270, 120662 (2021). 10.1016/j.biomaterials.2021.120662

23 Gunay, K. A. et al. Photo-expansion microscopy enables super-resolution imaging of cells embedded in 3D hydrogels. Nat Mater 22, 777–785 (2023). 10.1038/s41563-023-01558-5

24 Blatchley, M. R. et al. In Situ Super-Resolution Imaging of Organoids and Extracellular Matrix Interactions via Phototransfer by Allyl Sulfide Exchange-Expansion Microscopy (PhASE-ExM). Adv Mater 34, e2109252 (2022). 10.1002/adma.202109252

25 Bohin, N. et al. Rapid Crypt Cell Remodeling Regenerates the Intestinal Stem Cell Niche after Notch Inhibition. Stem Cell Reports 15, 156–170 (2020). 10.1016/j.stemcr.2020.05.010

26 Deng, F., Wu, Z., Zou, F., Wang, S. & Wang, X. The Hippo-YAP/TAZ Signaling Pathway in Intestinal Self-Renewal and Regeneration After Injury. Front Cell Dev Biol 10, 894737 (2022). 10.3389/fcell.2022.894737

27 Serra, D. et al. Self-organization and symmetry breaking in intestinal organoid development. Nature 569, 66–72 (2019). 10.1038/s41586-019-1146-y

28 Wang, T. C. et al. Matrix stiffness drives drop like nuclear deformation and lamin A/C tension-dependent YAP nuclear localization. Nat Commun 15, 10151 (2024). 10.1038/s41467-024-54577-4

29 Lima, J. T. & Ferreira, J. G. Mechanobiology of the nucleus during the G2-M transition. Nucleus 15, 2330947 (2024). 10.1080/19491034.2024.2330947

30 Wisniewski, E. O. et al. Dorsoventral polarity directs cell responses to migration track geometries. Sci Adv 6, eaba6505 (2020). 10.1126/sciadv.aba6505

31 Renkawitz, J. et al. Nuclear positioning facilitates amoeboid migration along the path of least resistance. Nature 568, 546–550 (2019). 10.1038/s41586-019-1087-5

32 Lomakin, A. J. et al. The nucleus acts as a ruler tailoring cell responses to spatial constraints. Science 370 (2020). 10.1126/science.aba2894

33 Venturini, V. et al. The nucleus measures shape changes for cellular proprioception to control dynamic cell behavior. Science 370 (2020). 10.1126/science.aba2644

34 Walker, C. J. et al. Nuclear mechanosensing drives chromatin remodelling in persistently activated fibroblasts. Nat Biomed Eng 5, 1485–1499 (2021). 10.1038/s41551-021-00709-w

35 Song, Y. et al. Transient nuclear deformation primes epigenetic state and promotes cell reprogramming. Nat Mater 21, 1191–1199 (2022). 10.1038/s41563-022-01312-3

36 Zhao, Z. et al. Organoids. Nat Rev Methods Primers 2 (2022). 10.1038/s43586-022-00174-y

37 Stanger, B. Z., Datar, R., Murtaugh, L. C. & Melton, D. A. Direct regulation of intestinal fate by Notch. Proc Natl Acad Sci U S A 102, 12443–12448 (2005). 10.1073/pnas.0505690102

38 Vooijs, M., Liu, Z. & Kopan, R. Notch: architect, landscaper, and guardian of the intestine. Gastroenterology 141, 448–459 (2011). 10.1053/j.gastro.2011.06.003

39 Lukonin, I. et al. Phenotypic landscape of intestinal organoid regeneration. Nature 586, 275–280 (2020). 10.1038/s41586-020-2776-9

40 Wester, R. A. et al. Retinoic acid signaling drives differentiation toward the absorptive lineage in colorectal cancer. iScience 24, 103444 (2021). 10.1016/j.isci.2021.103444

41 Dickinson, R. B. & Lele, T. P. Nuclear shapes are geometrically determined by the excess surface area of the nuclear lamina. Front Cell Dev Biol 11, 1058727 (2023). 10.3389/fcell.2023.1058727

42 Le, H. Q. et al. Mechanical regulation of transcription controls Polycomb-mediated gene silencing during lineage commitment. Nat Cell Biol 18, 864–875 (2016). 10.1038/ncb3387

43 Nava, M. M. et al. Heterochromatin-Driven Nuclear Softening Protects the Genome against Mechanical Stress-Induced Damage. Cell 181, 800–817 e822 (2020). 10.1016/j.cell.2020.03.052

44 Pashos, A. R. S. et al. H3K36 methylation regulates cell plasticity and regeneration in the intestinal epithelium. Nat Cell Biol 27, 202–217 (2025). 10.1038/s41556-024-01580-y

45 Burger, E. et al. Loss of Paneth Cell Autophagy Causes Acute Susceptibility to Toxoplasma gondii-Mediated Inflammation. Cell Host Microbe 23, 177–190 e174 (2018). 10.1016/j.chom.2018.01.001

46 Madisen, L. et al. A robust and high-throughput Cre reporting and characterization system for the whole mouse brain. Nat Neurosci 13, 133–140 (2010). 10.1038/nn.2467

47 Chakrabarti, R. et al. Notch ligand Dll1 mediates cross-talk between mammary stem cells and the macrophageal niche. Science 360 (2018). 10.1126/science.aan4153

48 Tikhonova, A. N. et al. The bone marrow microenvironment at single-cell resolution. Nature 569, 222–228 (2019). 10.1038/s41586-019-1104-8

49 DeForest, C. A. & Anseth, K. S. Cytocompatible click-based hydrogels with dynamically tunable properties through orthogonal photoconjugation and photocleavage reactions. Nat Chem 3, 925–931 (2011). 10.1038/nchem.1174

50 Bagby, S. et al. Development and Maintenance of a Preclinical Patient Derived Tumor Xenograft Model for the Investigation of Novel Anti-Cancer Therapies. J Vis Exp (2016). 10.3791/54393

51 Hawkins, H. J. et al. Examination of Wnt signaling as a therapeutic target for pancreatic ductal adenocarcinoma (PDAC) using a pancreatic tumor organoid library (PTOL). PLoS One 19, e0298808 (2024). 10.1371/journal.pone.0298808

52 Sato, T. et al. Long-term expansion of epithelial organoids from human colon, adenoma, adenocarcinoma, and Barrett’s epithelium. Gastroenterology 141, 1762–1772 (2011). 10.1053/j.gastro.2011.07.050

53 Seino, T. et al. Human Pancreatic Tumor Organoids Reveal Loss of Stem Cell Niche Factor Dependence during Disease Progression. Cell Stem Cell 22, 454–467 e456 (2018). 10.1016/j.stem.2017.12.009

54 Spence, J. R. et al. Directed differentiation of human pluripotent stem cells into intestinal tissue in vitro. Nature 470, 105–109 (2011). 10.1038/nature09691

55 McCracken, K. W., Howell, J. C., Wells, J. M. & Spence, J. R. Generating human intestinal tissue from pluripotent stem cells in vitro. Nat Protoc 6, 1920–1928 (2011). 10.1038/nprot.2011.410

56 Capeling, M. M. et al. Nonadhesive Alginate Hydrogels Support Growth of Pluripotent Stem Cell-Derived Intestinal Organoids. Stem Cell Reports 12, 381–394 (2019). 10.1016/j.stemcr.2018.12.001

