## Extended Figures for "Nuclei sense complex tissue shape and direct intestinal stem cell fate"

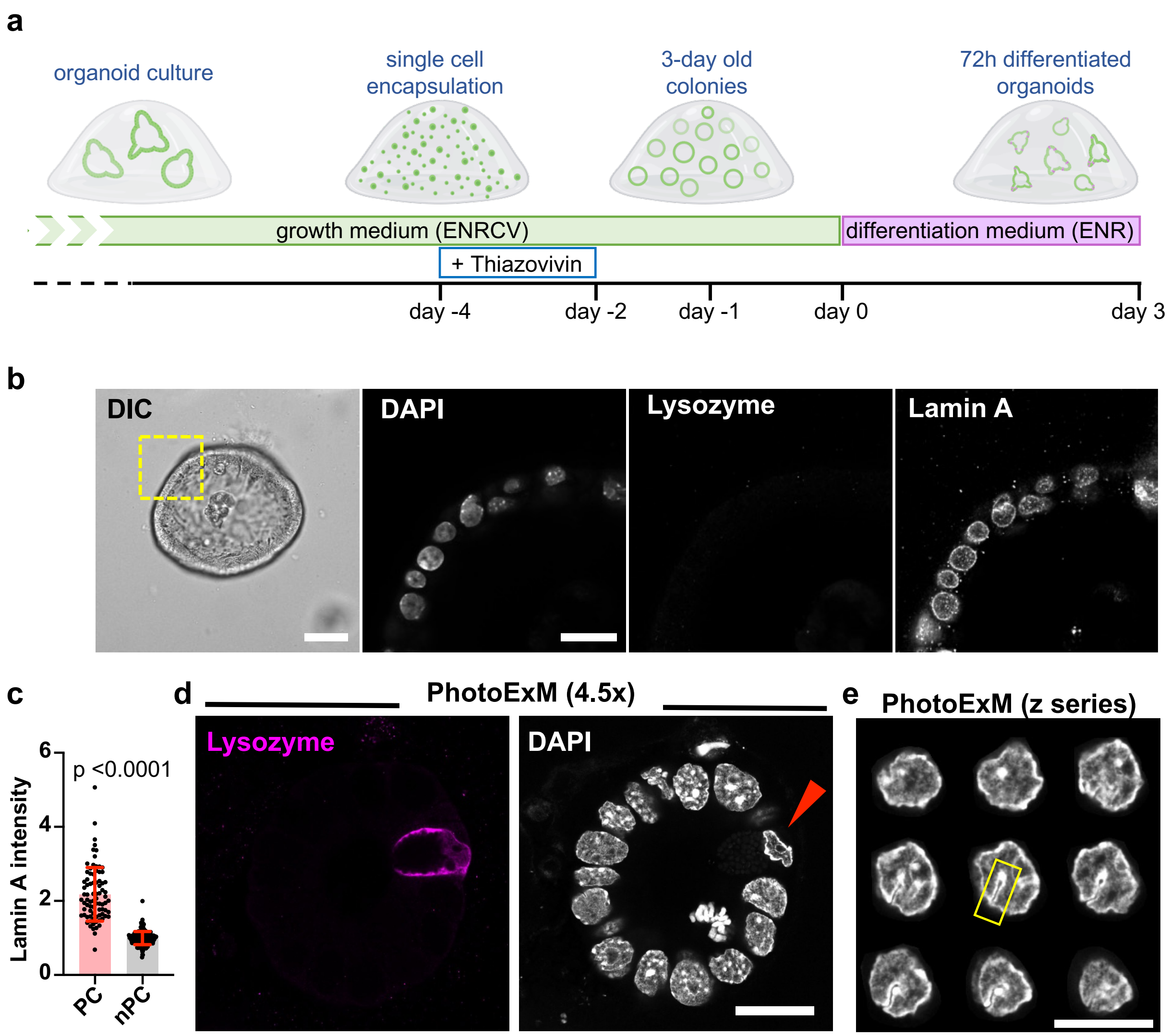


**Extended Figure 1: a**, Schematic of timeline and media conditions for experiments with organoids. **b**, *Left*, confocal DIC image of organoid kept in stem promoting ENRCV medium (scale bar = 50 μm). Confocal images of DAPI, lysozyme and lamin A show zoomed in area highlighted in yellow box in the DIC image (scale bar = 20 μm). Images are representative of 2 experiments. **c**, Lamin A intensities of NE in Paneth (PC) and non-Paneth cells (nPC) within *Defa4*$^{Cre}$/*Rosa26*$^{tdTomato}$ organoid crypts. Values normalized to average of nPC intensities in each image. Data are mean ± s.d. from 3 experiments. Statistical analysis was performed using Mann-Whitney U-test. **d**, Representative confocal images after 4.5X photo-expansion of differentiated organoid crypts immuno-stained for lysozyme showing deformed Paneth cell nucleus (red arrowhead). Images are representative of 2 experiments. Scale bar = 10 μm. **e**, Montage of representative z-sequence of DAPI stained Paneth cell nucleus from photo-expansion microscopy, showing furrow in DNA organization indicated with yellow box in the midplane image at center. Scale bar = 5 μm and z slices are 0.44 μm apart. Images are representative of 2 experiments.

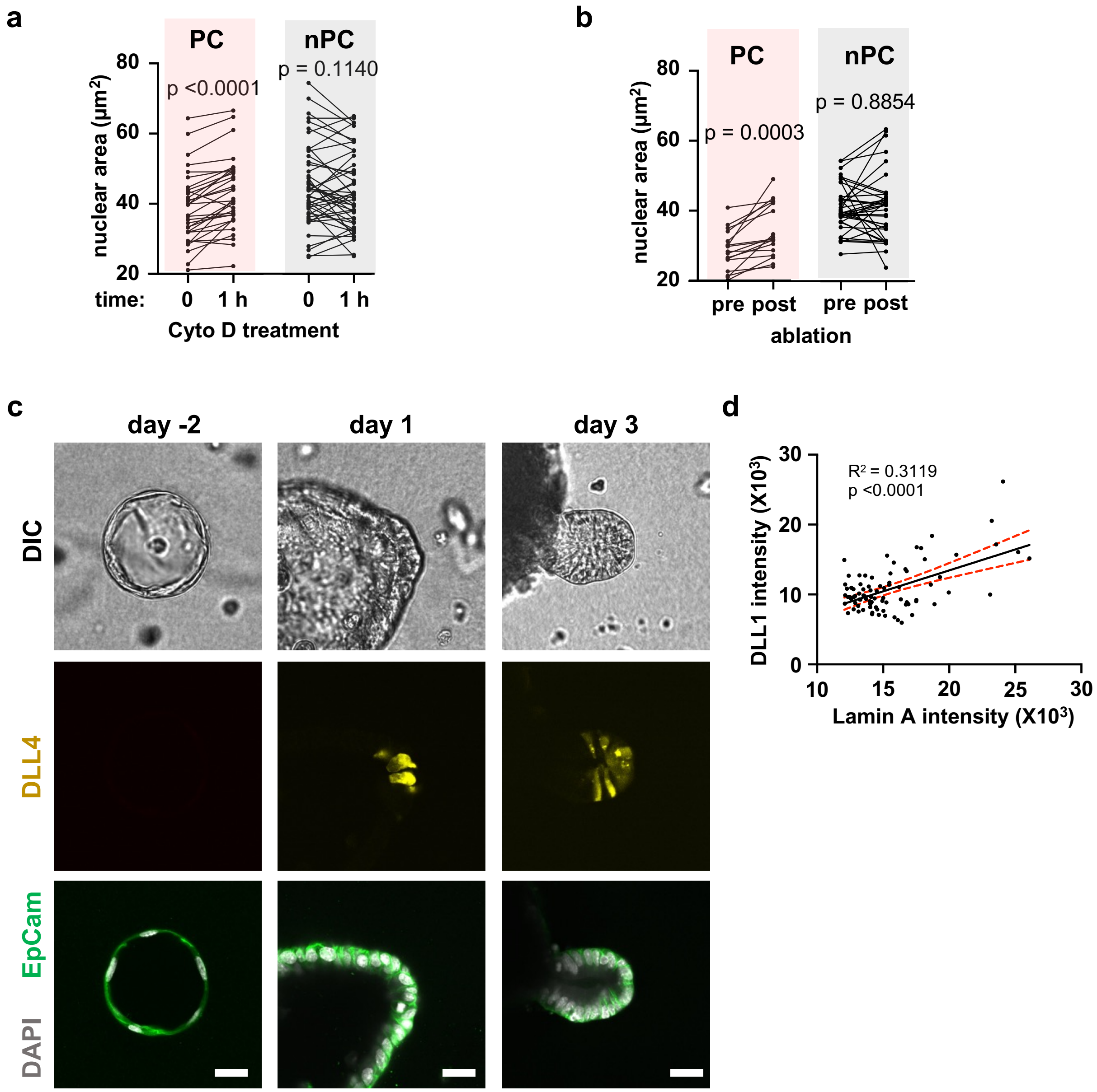


**Extended Figure 2: a**, **b**, Nuclear cross-sectional area of individual nucleus before and after 1 hour of Cytochalasin D treatment (a) and after photo-ablation (b). Data are from ≥ 3 experiments and paired t tests were used to derive statistics. **c**, Confocal images of DLL4-mCherry expressing organoids in Matrigel. Cells were seeded 4 days prior (day -4) to change in medium conditions from ENRCV to ENR (day 0). While there is no mCherry expression during their growth in ENRCV medium (day -2), mCherry-positive cells start appearing one day after ENR medium treatment (day 1). Images are representative of 3 experiments and scale bars = 20 μm. **d**, Correlation of lamin A and DLL1 immuno-staining in each cell, 3 days after differentiation in ENR medium. Solid line indicates linear regression and dashed red lines define the 95% confidence intervals from 2 experiments.

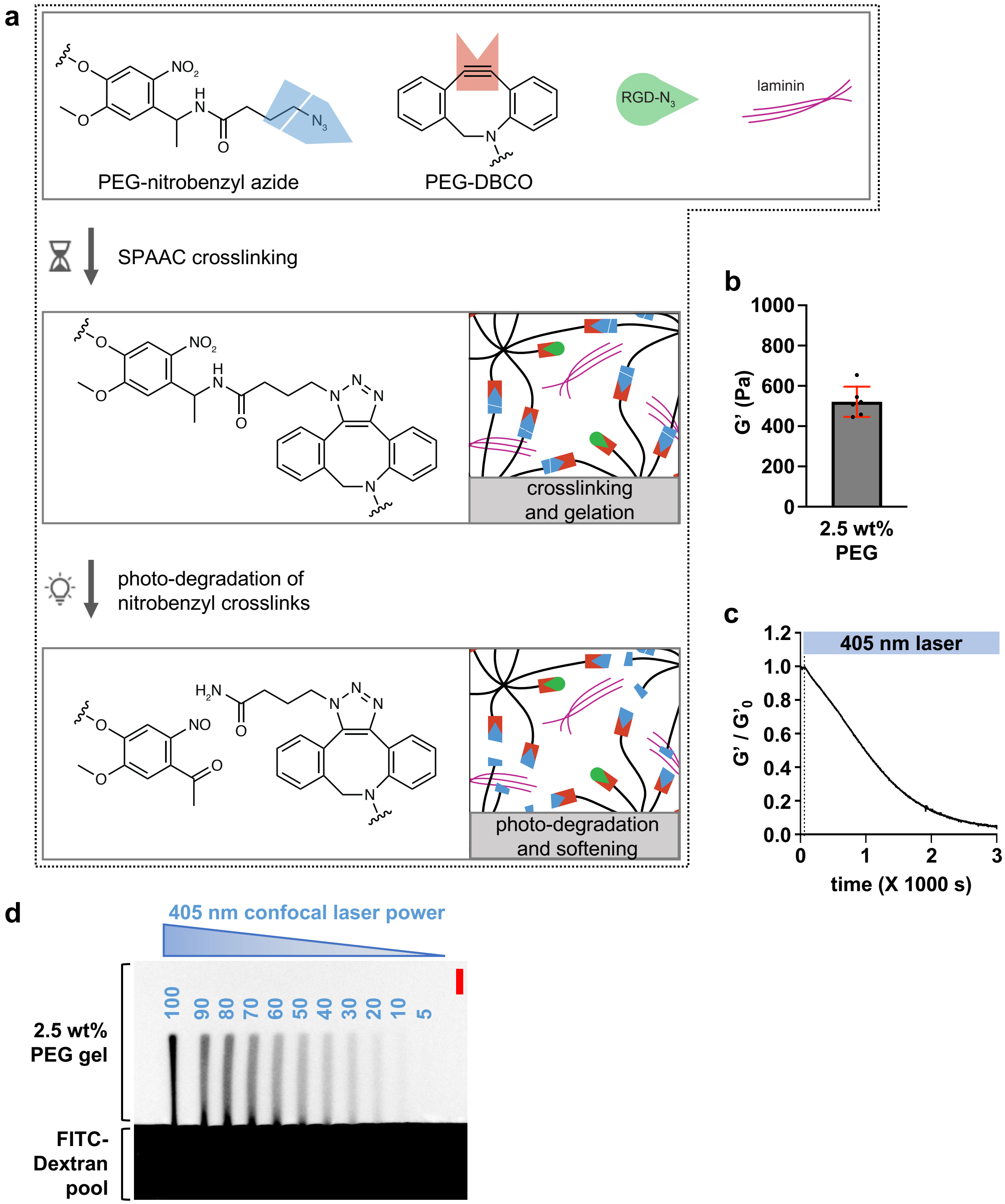


**Extended Figure 3: a,** Schematics of PEG based hydrogel formulation, crosslinking, and photo-degradation. **b**, Hydrogels fabricated with 2.5 weight percent of total polymer achieved an equilibrated mean shear storage modulus of 520 Pa after overnight incubation in PBS. Dots indicate the shear storage modulus of each gel and lines indicate s.d. **c**, Normalized shear storage modulus of the photo-degradable gel which softened upon illumination with 50 mW $cm^{-2}$ 405 nm laser started after 60 s, as indicated by the blue bar. Trace is representative of 3 gels and data normalized to the initial storage modulus ($G'_0$). **d**, Confocal image of 2.5 wight percent PEG gel incubated with a $2X10^6$ Da FITC Dextran solution (FITC signal indicated in black). FITC Dextran solution was present at the edge of the gel and had entered 40 μm wide rectangular regions irradiated with one photon 405 nm confocal laser operated at indicated power levels ranging from 5% to 100%. Scale bar = 100 μm.

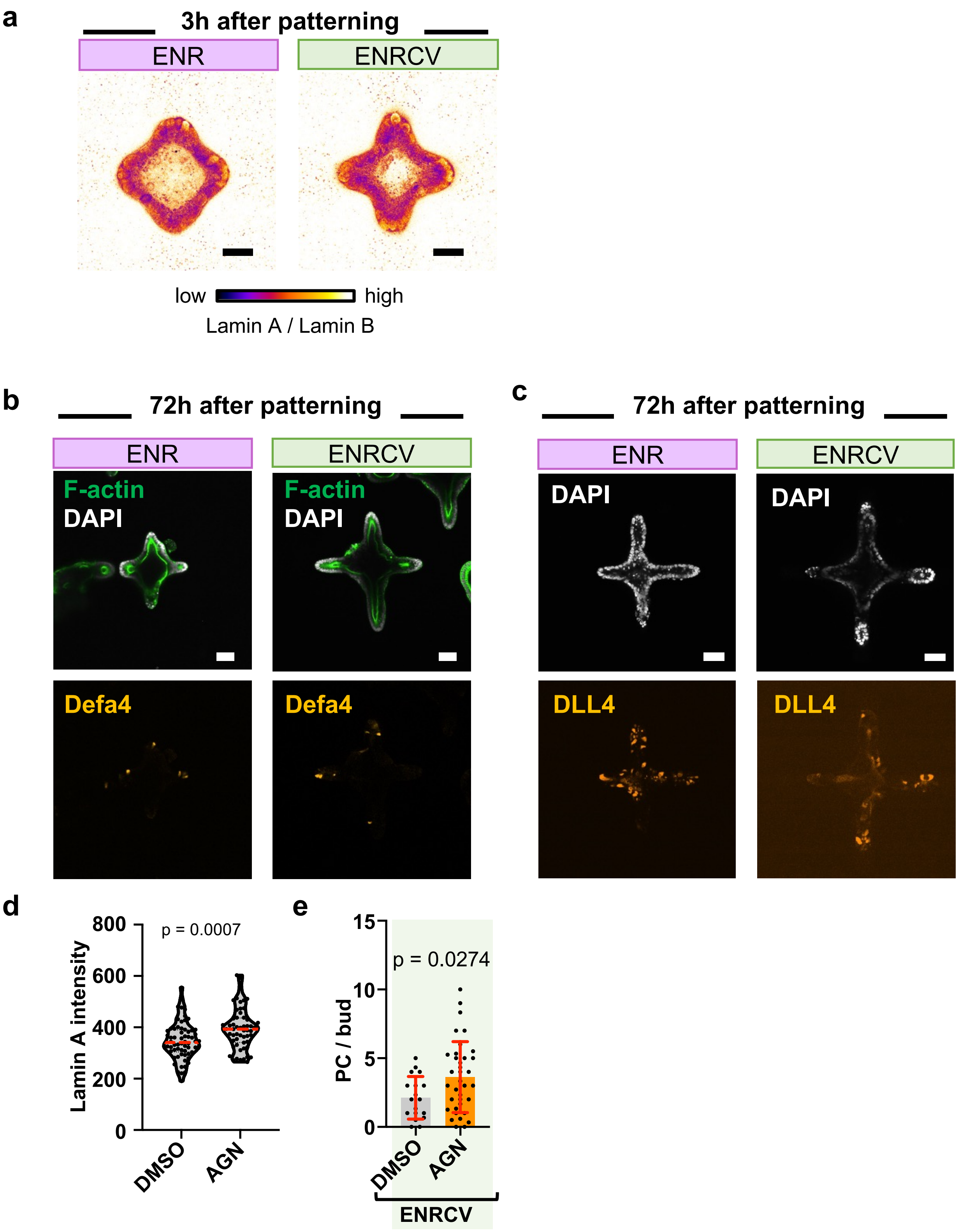
a
3h after patterning
ENR
ENRCV
low
high
Lamin A / Lamin B
b
72h after patterning
ENR
ENRCV
F-actin
DAPI
Defa4
c
72h after patterning
ENR
ENRCV
DAPI
DLL4
d
p = 0.0007
Lamin A intensity
800
600
400
200
0
DMSO
AGN
e
p = 0.0274
PC / bud
15
10
5
0
DMSO
AGN
ENRCV

**Extended Figure 4:** **a**, Immuno-stained confocal images of organoids 3 hours after patterning and incubation in ENR or ENRCV medium. Heatmaps show pixel-wise ratio of lamin A over lamin B immunofluorescence. Images are representative of 3 experiments. Scale bars = 20 µm. **b**, **c**, Confocal images of photo-patterned *Defa4*$^{Cre}$/*Rosa26*$^{tdTomato}$ (b) or *Dll4-mCherry* (c) organoids after 3 days of incubation with ENR or ENRCV medium. Images are representative of 3 experiments**.** Scale bars = 50 µm. **d,** Lamin A intensity in immuno-stained organoids treated with AGN for 3 days. Lines indicate the medians and data are from 2 experiments. **e,** Average number of Paneth cells in each bud for organoids treated with AGN or vehicle control. Data are mean ± s.d. from 3 experiments. Statistical analysis was performed using unpaired t test (in **d** and **e**).

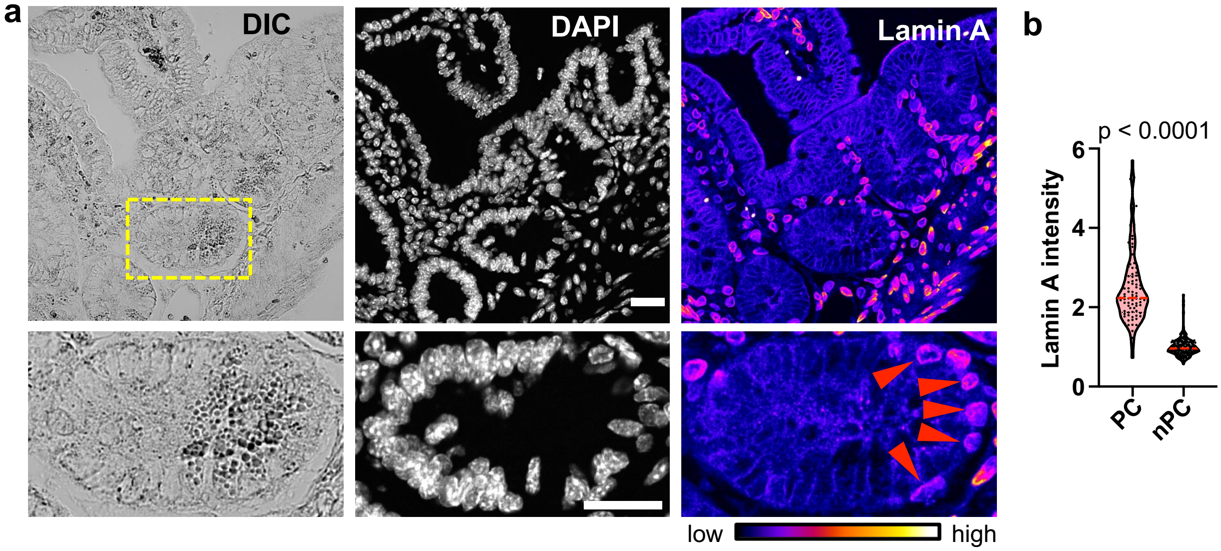


**Extended Figure 5:** **a**, Confocal images of intestinal section from wild-type mouse stained for DAPI and lamin A. Bottom row shows zoomed in view of the region indicated by yellow box. Heatmaps show level of lamin A and arrowheads indicate high lamin A intensity in the nuclei of Paneth cells. Paneth cells were identified by the presence of detectable lysozyme granules in DIC images. Images are representative of 2 mice and scale bars = 20 µm. **b**, Intensities of lamin A in Paneth and non-Paneth cell nuclei within tissue sections. Values normalized to average of nPC intensities in each image. Data are from 2 mice; the dashed red lines indicate the median and dotted black lines indicate the quartiles. Statistical analysis was performed using the Mann-Whitney U-test.
